# Exploring rare cellular activity in more than one million cells by a trans-scale-scope

**DOI:** 10.1101/2020.06.29.179044

**Authors:** T. Ichimura, T. Kakizuka, K. Horikawa, K. Seiriki, A. Kasai, H. Hashimoto, K. Fujita, T. M. Watanabe, T. Nagai

## Abstract

In many phenomena of biological systems, not a majority, but a minority of cells act on the entire multicellular system causing drastic changes in the system properties. To understand the mechanisms underlying such phenomena, it is essential to observe the spatiotemporal dynamics of a huge population of cells at sub-cellular resolution, which is difficult with conventional tools such as microscopy and flow cytometry. Here, we describe an imaging system named AMATERAS that enables optical imaging with an over-one-centimeter field-of-view and a-few-micrometer spatial resolution. This trans-scale-scope has a simple configuration, composed of a low-power lens for machine vision and a hundred-megapixel image sensor. We demonstrated its high cell-throughput, capable of simultaneously observing more than one million cells. We applied it to dynamic imaging of calcium ions in HeLa cells and cyclic-adenosine-monophosphate in *Dictyostelium discoideum*, and successfully detected less than 0.01% of rare cells and observed multicellular events induced by these cells.

## Main text

One of the challenges faced by cell biology research is the understanding of the systems in which rare cells in a large population play an important role in determining the fate of the entire multicellular system. Typically, in conventional methodology, cells or cell-type comprising the majority group are focused upon, and rare cells are often excluded as noise or outliers. Hence, the function of such rare cells has not been well studied. To understand the biological importance of rare cells, it is necessary to measure and analyze all individual cells in a multicellular system. Such issues are common in various research areas of medicine and biology, including developmental biology,^1^ neuroscience,^2,3^ oncology,^4^ and immunology.^5^ To tackle this challenge, it is essential to develop an optical microscope system for single-cell imaging within macroscale dynamics in a wide field-of-view (FOV), which enables dynamic observation of a huge number of cells at the same time.

This multiple scale-hierarchy observation system can be termed as “trans-scale-scope”. Since the spatial resolution and magnification of optical microscope are correlated owing to the structure of the imaging lens system, a high (or low)-resolution imaging system has a high (or low) magnification. This fundamental dilemma makes it difficult to expand the FOV while maintaining sub-cellular spatial resolution. To achieve this, an increase in the number of sampling points, *i.e.*, the pixels in an image sensor in wide-field imaging and scan points in laser-scan imaging, is indispensable. Specifically, 4-megapixel scientific complementary metal oxide semiconductor (CMOS) sensors are widely used in recent cell biology. If wide-field imaging is performed under low magnification conditions, such as 100 pixels (10 × 10) per cell, the number of cells that can be observed simultaneously is not more than 4 × 10^4^.

To overcome this limitation, we propose to build an observation system for high cell-throughput using a CMOS sensor with more than hundred megapixels for machine vision, which is not usually used in biological microscopes. As a proof-of-concept, we adopted a commercially available CMOS image sensor, 120MXSM, developed by CANON Inc., which is one of the image-sensors with the largest number of pixels on the market. The CMOS chip has 13264 × 9180 effective pixels and exhibits a 2.2 µm pixel size and 29.2 mm × 20.2 mm chip size. The small pixel size eliminates the need for large magnification in studies requiring spatial resolution at the sub-cellular level. The size of this CMOS chip is 35 mm diagonal, which is more than the field-number (*FN*) of a normal biological microscope objective, typically 22–25 mm. To make the best use of the entire sensor area, we introduced a low-power lens with a large *FN*. Specifically, we adopted a commercially available telecentric 2× lens for machine vision. This simple configuration enables cellular imaging with sub-cellular resolution in a vast FOV of 14.6 × 10.1 mm^2^, which can achieve cell throughput of more than one million cells in a single shot.

Compared to trans-scale-scopes recently reported by other groups,^6–10^ our imaging instrument has been designed with a higher emphasis on the FOV. Although the spatial resolution is not high enough to resolve the intracellular molecular distribution, the FOV and cell throughput are unrivaled by the others. In addition to this high-throughput feature, the optical configuration of this method is simple and low-cost, making it suitable for wide dissemination in laboratories of biology, pharmacology and medicine. Thus, it is a promising tool to open-up the scientific field on the function of rare cells.

We named our trans-scale-scope AMATERAS (a multi-scale/modal analytical tool for every rare activity in singularity). Although we aim to eventually establish AMATERAS as a multimodal measurement instrument, the present version (AMATERAS1.0) can be employed only for a single modality (optical imaging). We demonstrated that AMATERAS1.0 has the potential to observe more than one million cells simultaneously within one second. We then employed the AMATERAS1.0 for calcium ion (Ca^2+^) imaging of HeLa cells, and cyclic-adenosine-monophosphate (cAMP) imaging in *Dictyostelium discoidium* (*D. discoidium*) cells. We demonstrate a successful observation of rare events and cells.

## Results

### Configuration and performance of AMATERAS1.0

We selected a wide-field imaging configuration for AMATERAS1.0 (Fig. 1A), rather than laser scan imaging, and used a single CMOS chip as it facilitates the recording of the entire FOV. We used a camera (VCC-120CXP1M, CIS, Tokyo, Japan) equipped with the 120-megapixel image-sensor mentioned in the Introduction. Imaging lenses for this image-sensor (35 mm diagonal) at such magnifications are not in the line-up of the objective lens of biological microscopes but are available in those used for machine vision in which telecentric lenses are utilized, owing to the requirements of the measurements with high linearity and less distortion. We selected a 2× telecentric macro-lens for a full-size image sensor (*FN* = 44 mm, LSTL20H-F, Myutron, Japan) owing to its relatively high numerical aperture (*NA* = 0.12 at the object side). The combination of the above image sensor and the lens allows for imaging with a FOV of 14.6 × 10.1 mm^2^ (17.8 mm diagonal) and a spatial resolution better than 2.5 µm in the visible wavelength region. This FOV is wide enough to observe one million confluent eukaryotic cells in a single shot.

**Fig. 1.**
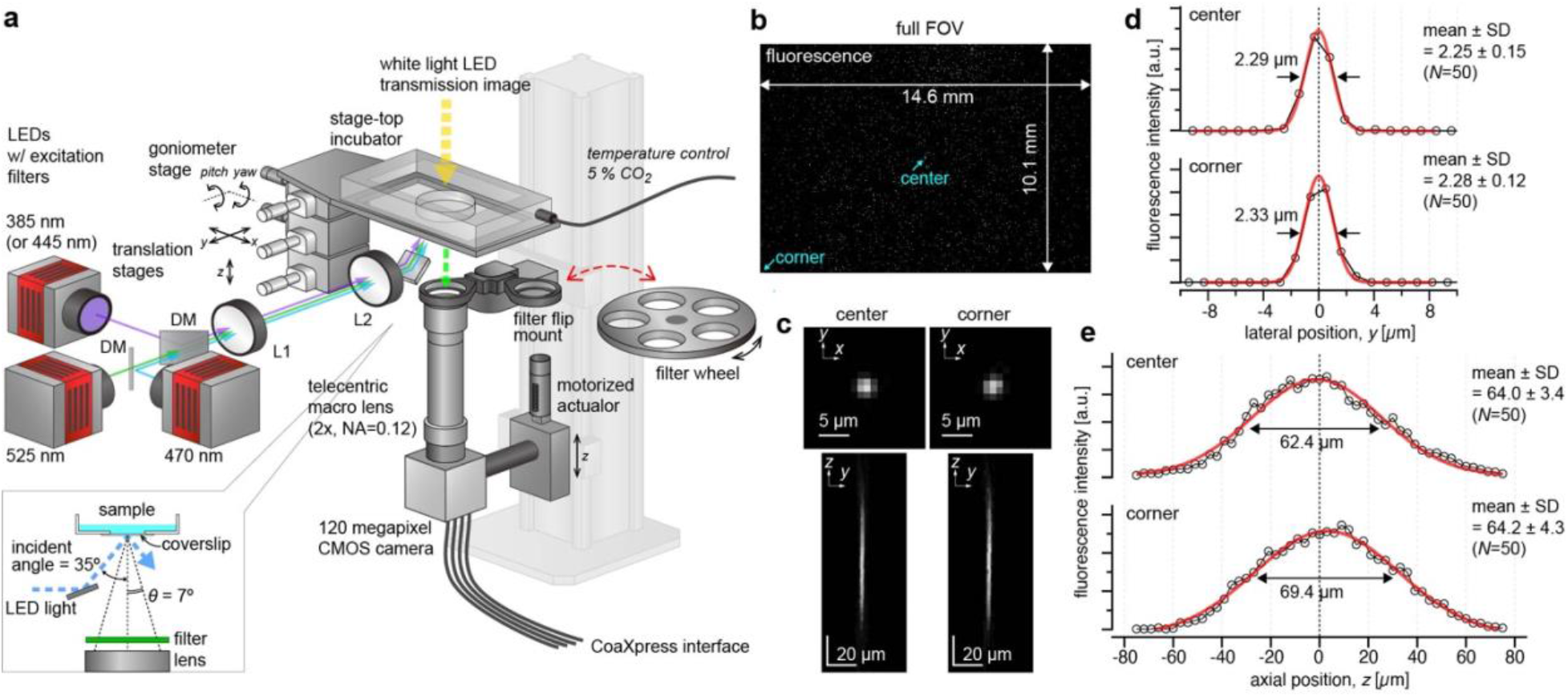
Configuration and performance of the trans-scale imaging system AMATERAS1.0. (a) Schematic representation of the system configuration. (b-e) Evaluation of the imaging performance with fluorescent beads of 200 nm in diameter dispersed on a glass-bottom dish. The central wavelength of LED-excitation was 470 nm, and the emission peak wavelength was 520 nm. (b) An image captured with full FOV. (c) Representative images of individual beads in a transverse plane (*xy* plane) and a longitudinal plane (*yz* plane) at the center and the corner of the FOV (indicated by light-blue arrows in b), which can be regarded as PSF. The *yz* plane image is a longitudinal cross-section of an image-stack obtained by scanning the sample-lens distance in the *z*-direction. (d) Line profiles depicting the fluorescence intensities on the *y*-axis at the in-focus plane (c, top), shown with black circles. (e) Line profiles depicting the fluorescence intensities on the *z*-axis penetrating the center of the PSF (c, bottom), shown with black circles. The red lines in (d-e) represent a Gauss function curve fit to the experimental data.

The entire optical system of AMATERAS1.0 is amazingly simple, consisting of the aforementioned camera-lens pair and the illumination light source with its relay optics (Fig. 1a, See Methods for more detail). High-brightness LEDs (>3 W) with four different colors (SOLIS series, Thorlabs, Newton, NJ) were used for fluorescence excitation which were combined at dichroic beam-combiners. Taking advantage of the long working distance of the imaging lens (∼110 mm), the excitation light was introduced obliquely from the bottom of the sample at an incident angle of 35° greater than the half angular aperture (α ∼ 7°) of the imaging lens. This illumination geometry allows us to prevent directly reflected light from entering the imaging lens; hence, fluorescence imaging can be performed with only a single emission filter placed at the entrance of the imaging lens and does not require dichroic mirror. In multi-color fluorescence imaging, the emission filters were switched by the motorized filter wheel. Bright-field imaging can also be performed with another white-light LED source placed at the top of the sample.

To evaluate the spatial resolution, we experimentally recorded the point-spread-function (PSF) by imaging green fluorescent beads with 200 nm diameter (wavelength ∼520 nm). Figures 1b-c shows (b) a full FOV image of the beads and (c) representative images of single beads in transverse (*xy*) and longitudinal (*yz*) planes at the center and corner of the FOV. The lateral spatial resolution and axial depth-of-focus were evaluated using full-width-at-half-maximum (FWHM) of the Gaussian function fit to the PSF. The FWHM in the transverse plane was 2.25 µm and 2.28 µm at the center and corner, respectively (Fig. 1d), but the difference was not statistically significant (*p* = 0.24, Student’s *t*-test with *N* = 50). The experimental FWHM values were close to the theoretical value, 2.17 µm (∼ 0.51*λ*/*NA*, *NA* = 0.12, *λ* = 520 nm). The depth-of-focus was ∼64 µm (Fig. 1e) with no significant difference between the center and corner (*p* = 0.85), which is consistent with a numerically calculated value of 60.6 µm (Fig. S1a-c, Note 1). There is a concern on the presence of glass plates (170-µm-thick coverslip of sample dish and a 2-mm-thick emission filter) between the sample and lens, which causes a spherical aberration. However, because of the low *NA*, the PSF broadening by spherical aberration was almost negligible both in the lateral and axial directions, as clarified by a numerical calculation (Fig. S1c-e, Note 1). These results indicate that the spatial resolution equivalent to the theoretical one can be achieved in the entire FOV.

We compared the imaging property of AMATERAS1.0 with a configuration in which its lens or camera was replaced by a conventional one in terms of the pixel resolution and FOV (Fig. S2, Note 2). We also compared it with trans-scale imaging techniques developed by other groups in terms of spatial resolution and FOV (Fig. S3, Note 3). These comparisons clarified the advantages and features of our system.

### Brain slice imaging

To demonstrate trans-scale observation of a biological specimen, we performed multi-color imaging of a centimeter-wide mouse brain section. Two types of fluorescent proteins, tdTomato^11^ and EGFP, were expressed in excitatory projection neurons and inhibitory interneurons, respectively. In addition, the nuclei were stained with Hoechst 33342. Figure 2 shows an RGB image reflecting the fluorescence intensities of tdTomato (red), EGFP (green), and Hoechst (blue). The left panel exhibits the full FOV in which the whole brain is observed within the single FOV. Any specific region can be zoomed in to see the local distribution of cells at a single-cell resolution as if it were a Google map. For instance, two regions of the brain indicated by the light-blue squares (*A*: cerebral cortex, *B*: hippocampus) were digitally magnified 5-fold, as shown in the middle top and bottom panels. The light-blue square region in the 5× images was further magnified by 5-fold in the right panels, where the distribution of single neuronal cells and even neurite was clearly observed.

**Fig. 2.**
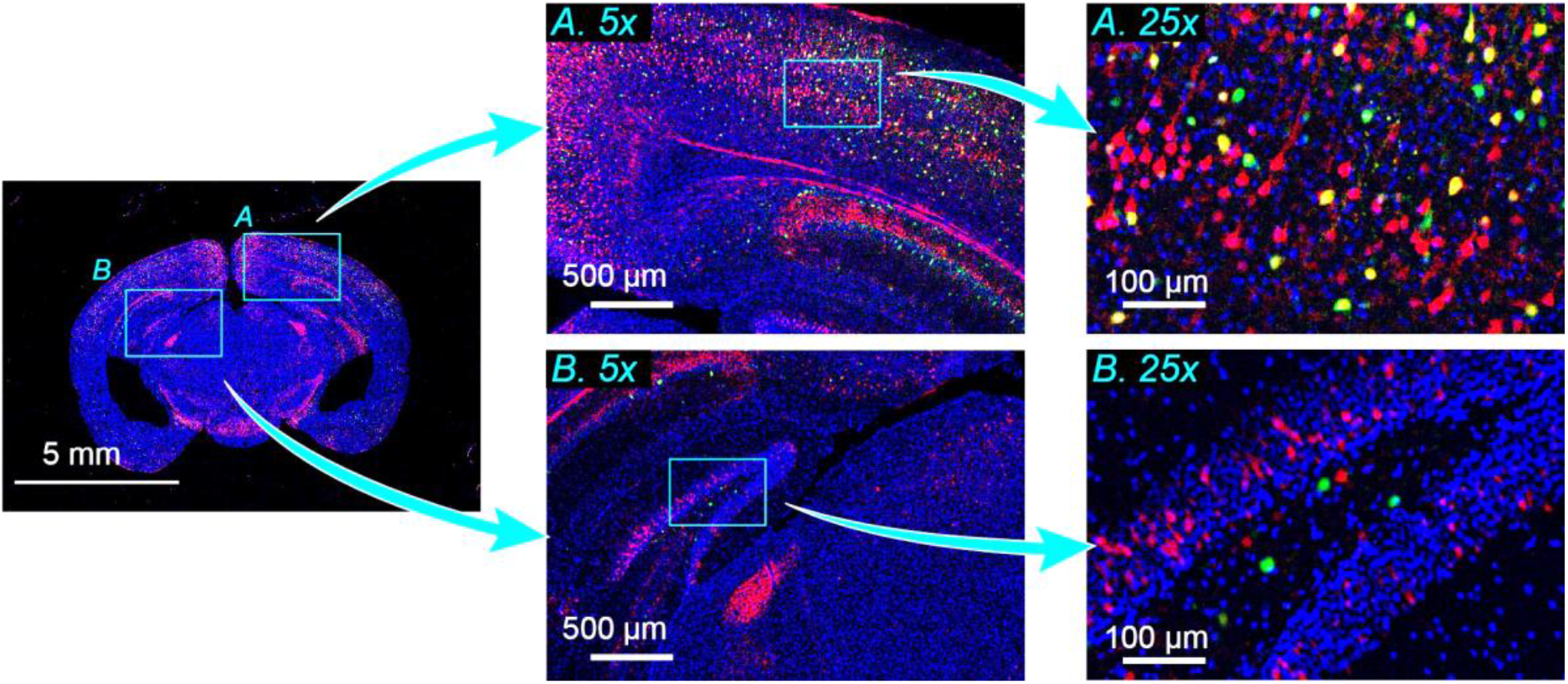
Imaging of the mouse brain. Multi-color image of a mouse brain slice with a thickness of 25 µm. Two regions indicated by light-blue squares, namely, cerebral cortex (*A*) and hippocampus (*B*), in the whole brain image (left) are magnified by 5-fold (middle). The local regions of light-blue squares in the 5x images are further magnified by 5-fold (right). Red, green, and blue represent the fluorescence intensity of tdTomato expressed in excitatory projection neurons, EGFP expressed in inhibitory interneurons, and Hoechst 33342 attached to nuclear DNA, respectively. The three color-channels were excited by the use of three LED wavelengths (center wavelengths: 525 nm, 470 nm, 385 nm) with an exposure time of two seconds for each channel.

These results also demonstrated the potential of AMATERAS1.0 in expediting whole-brain imaging. In contrast to previous technologies,^12,13^ the present method is capable of imaging a section with a size of over one centimeter in a single shot without the need for tiling or scanning, leading to a considerable reduction in the total imaging time required to observe a three-dimensional whole brain.^13^

### Single shot detection of more than million cells

We next demonstrate the high cell-throughput of the AMATERAS1.0 by observation of epithelial cells in the confluent state. The nuclei of the fixed Madin-Darby Canine Kidney (MDCK) cells were stained with a nucleus-staining green fluorescent chemical probe, NucleoSeeing (Funakoshi, Tokyo, Japan).^14^ The bright-field transmission images and fluorescence images in Fig. 3a verified that the individual cells could be imaged separately over the entire FOV. The number of cells in the FOV was determined to be approximately 1.19 × 10^6^, over one million. This throughput is much higher than that observed using normal microscopes. The mean area occupied by single cells was 123.8 µm^2^ (equivalent to 102.3 pixels).

**Fig. 3.**
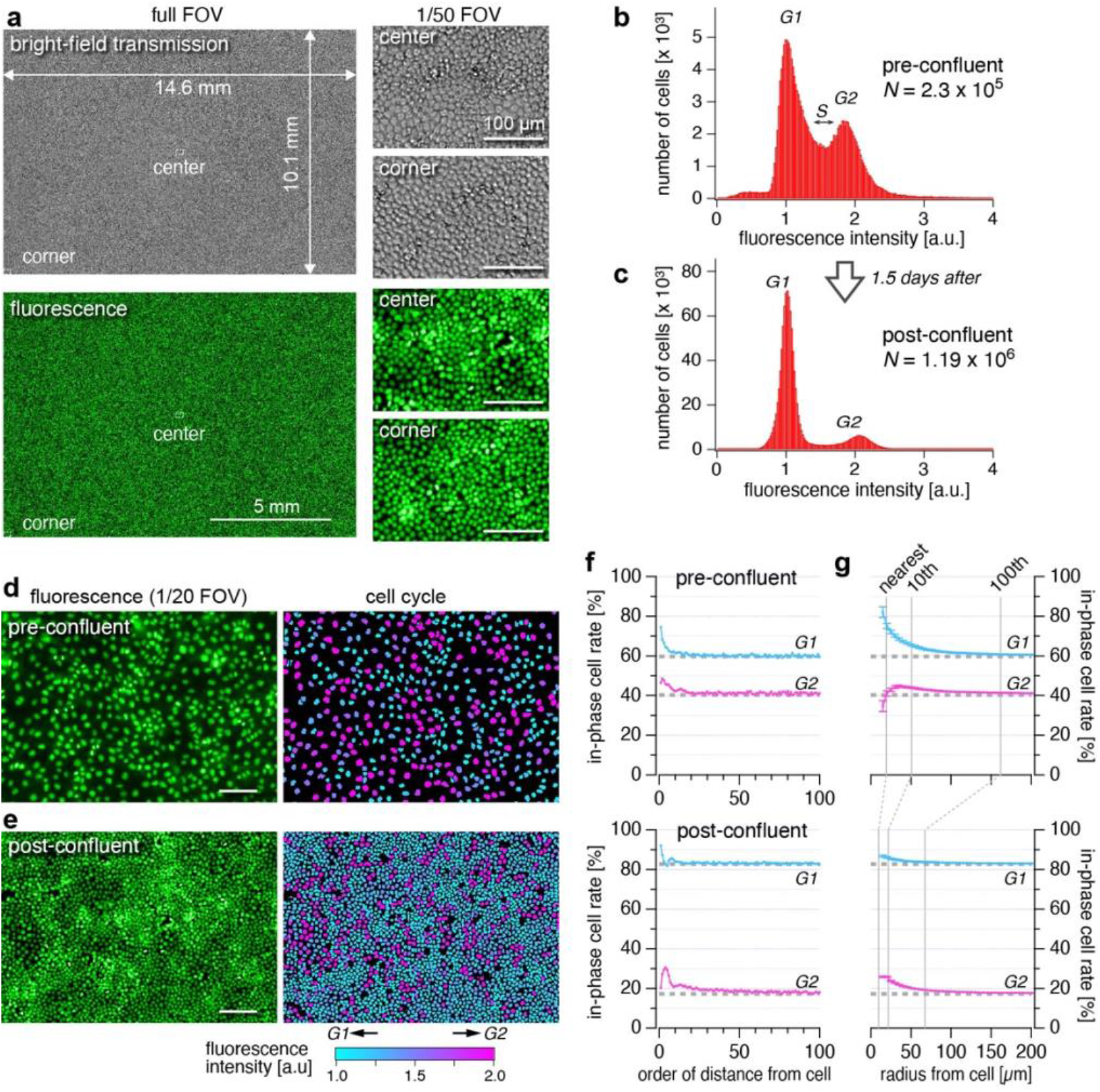
Single shot detection and analysis of more than one million cells. Imaging of the cultured MDCK cells that were fixed with paraformaldehyde and stained with NucleoSeeing was performed with an excitation LED wavelength of 470 nm. (a) Full FOV image and closeup images of the area covering 1/50 of the FOV region at the center and corner, obtained with the bright-field transmission (top three) and fluorescence (bottom three) modes. (b-c) Histograms of fluorescence intensity of cells with two different cell densities (b: pre-confluent, c: post-confluent), which correspond to the populations of cells in the *G1* and *G2* phases. The fluorescence intensity was normalized by the peak position for the *G1* phase. (d-e) Closeup views of areas comprising 1/20 of the FOV region obtained at the pre-confluent condition (d) and the post-confluent condition (e). In the right panels, cellular nuclei are painted with colors indicating fluorescence intensities of the cells, with each color corresponding to different phases of the cell cycle, *i.e.*, cyan and magenta represent the *G1* and *G2* phases, respectively. Scale bars: 100 µm. (f-g) Spatial analysis of the cells in the two phases. (f) In-phase cell rate of neighboring cells as a function of order of cell-cell distance (*G1*: cyan, *G2*: magenta). (g) In-phase cell rate among the cells in the neighboring area was plotted as a function of the radius of the area (*G1*: cyan, *G2*: magenta). The error bars represent standard errors. The dashed grey lines in f and g represent the expected values for random distribution.

We extracted the feature parameters of all individual cells in the entire FOV by using a machine learning software for cell image analysis (AIVIA 9.0, DRVision Technologies LLC, Bellevue, WA). We plotted fluorescence intensities of nuclei for the populations of MDCK cells at two different cell-densities, pre-confluence and post-confluence (1.5 days after pre-confluence) with estimated number of cells in the FOV 2.3 × 10^5^ and 1.19 × 10^6^, respectively (Fig. 3b-c). The bimodal distribution in both the two states corresponds to the populations of amount of DNA in individual cells, which can be attributed to populations of cells in the *G1* and *G2* phases. Cells in the region between the two peaks are considered to be in the *S* phase. A significant difference in the relative height of the two peaks for the *G1* and *G2* phases was found between Fig. 3b and 3c. This can be explained by the fact that *G1*-*S* phase transition rate is suppressed by contact inhibition in the highly confluent state,^15^ and is consistent with the previous studies using flow-cytometry.^16^

In contrast to flow-cytometry in cell cycle analysis,^17,18^ the present method provides information on the cell phase population along with their spatial distribution. In Fig. 3d-e, the cell nuclei in the original image are painted with colors indicating the fluorescence intensities of the cells, where each color represents the different phases of the cells. This color map suggests that under the two cell densities, cells in the same phase (in-phase cells) tend to be located closely. In order to quantitatively evaluate the non-randomness of the spatial distribution, we calculated two measures of similarity of neighboring cells, in-phase cell rate of neighboring cells as a function of order of distance and of radius of neighboring area (See Method and Fig. S4), as plotted in Fig. 3f-g, respectively. Figure 3f obviously showed that, in both the conditions, the in-phase cell rates for closer cells were significantly higher than the expected values for random distribution, and that for distal cells asymptotically approaches to the expected values. The second analysis (Fig. 3g) also verified the common tendency for cells in the same phase to cluster together, and furthermore provided additional information on the spatial scale of the clusters, which was found to be different between the two cell densities. By exponential fit, the mean sizes of the clusters were found to be about 60 µm and 30 µm for the pre- and post-confluent conditions, respectively. The clustering can be interpreted as cells originating from the same cell are still in phase over a few generations.^19^ The mechanical interactions of neighboring cells to synchronize their phases may also be involved.^20^

### Rare event detection by calcium imaging in HeLa cells

To demonstrate the possibility of detecting rare cells in the dynamics of living cells, we applied the trans-scale-scope to the imaging of Ca^2+^ concentration ([Ca^2+^]) dynamics in HeLa cells. For visualization of Ca^2+^ concentration, we used a fluorescent Ca^2+^ indicator, Yellow Cameleon 3.60 (YC3.60), which is based on a [Ca^2+^]-dependent induction of Förster resonance energy transfer (FRET) between an enhanced cyan-fluorescent protein, ECFP, and a yellow fluorescent protein, Venus. ^21^ Basically, this fluorescent probe enables quantitative imaging by the ratio of the fluorescence intensity of two wavelength channels, but here, we observed only the FRET intensity at the wavelength of Venus. This results in a loss of quantitative capability but no loss of imaging speed.

Figures 4a-c show the distribution of the signal intensity in the FRET channel in the full FOV (a), a 16× digitally magnified image of the dashed square region (b), and a bright-field image of the same region (c). The number of cells in the FOV was estimated to be 1.2 × 10^5^. Although it is well known that the majority of cells typically increase [Ca^2+^] upon stimulation with ligands such as histamine (Fig. S5, Video 1, Note 4), here, observations were made under conditions where histamine was not added. Nevertheless, time-lapse imaging at 5-second intervals revealed that a tiny fraction of cells exhibited spontaneous transient increases in [Ca^2+^] (Video 2).^22^ By an analysis of temporal profile at all cells, we searched for all the unique cells that spontaneously increase their [Ca^2+^]. The number of calcium-pulsing cells was found to be 394 cells in 200 frames (1,000 s), corresponding to approximately 2.0 cells per frame (0.0017% of 1.2 × 10^5^ cells). The rest of more than 99 % of cells did not exhibit significant [Ca^2+^] oscillation. The temporal profiles of their [Ca^2+^] of all the 394 cells were arranged in Fig. 4d, and five representative profiles are shown in Fig. 4e including one for a non-pulsing cell (*B*). The positions of the pulsing cells are marked on the full FOV image with the red-white-blue color representing the pulsing time in Fig. 4f, and the time course of the number of pulsing cells in successive six frames is shown in Fig. 4g-top.

**Fig. 4.**
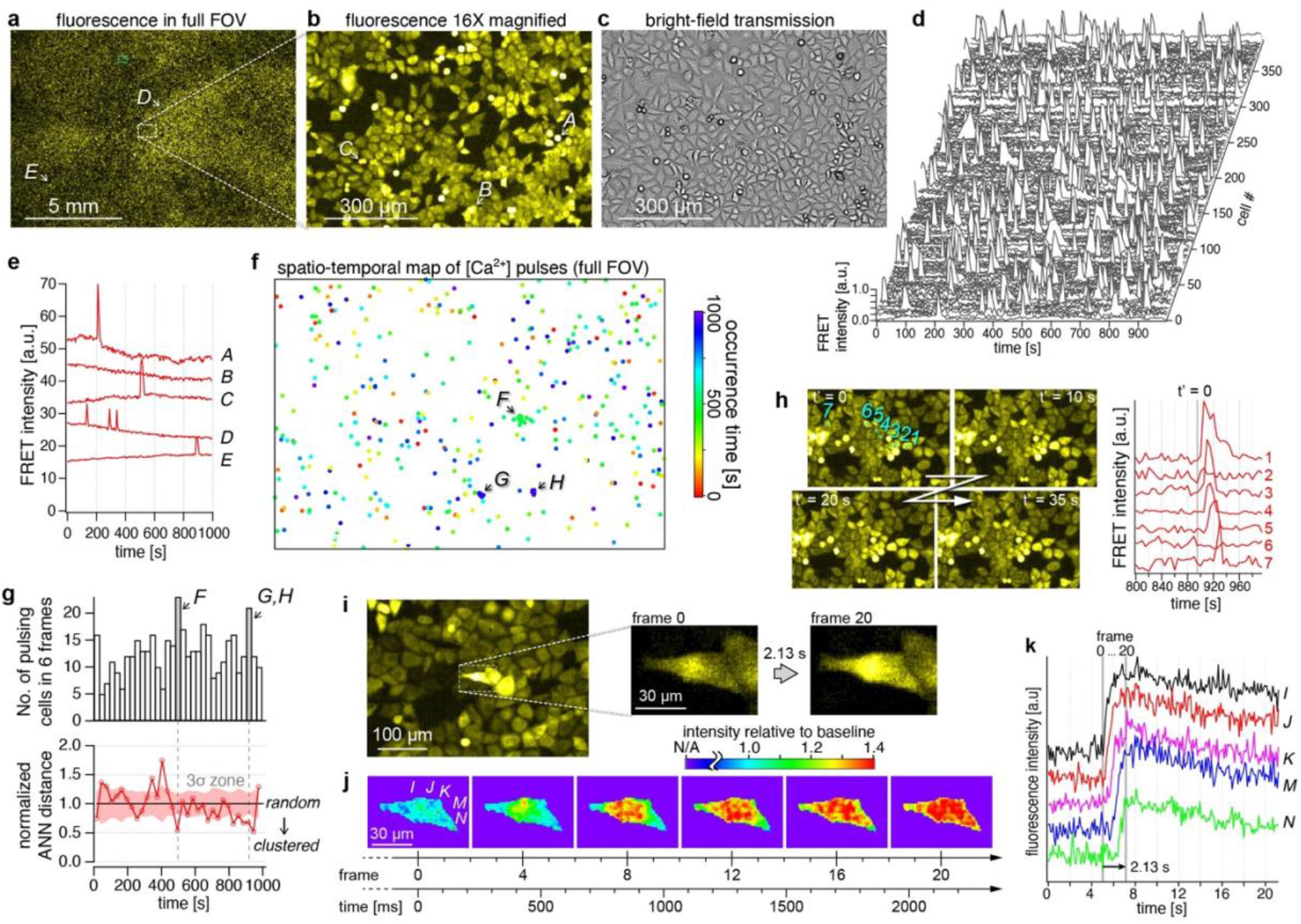
Calcium ion imaging of HeLa-YC3.60 cells. (a-b) Fluorescence intensity image of the FRET channel in (a) the full FOV, and (b) a 16× magnified image of the dashed square region indicated in (a). (c) Bright-field image of the region described in (b). (d) Temporal profiles of FRET channel intensity of 394 detected cells exhibiting spontaneous calcium pulse. (e) Temporal profiles of FRET channel intensity of five cells indicated with alphabets *A*-*E* in (a) and (b), including those exhibiting 1 and 3 peaks of spontaneous [Ca^2+^] pulses (*A*, *C-E*) and one with no peak (*B*). Images were acquired at 5-second intervals. (f) A plot of all cell positions detected in the 1,000-second measurement time. The occurrence time is indicated with red-white-blue color. (g) Time variation of the number of pulsing cells in the full FOV (top) and normalized average nearest neighbor (ANN) distance (bottom). The time bin width of the histogram is 6 frames (= 30 s). The light-red shade represents three times the width of the standard deviation. (h) Four image frames showing the [Ca^2+^] propagation at the location *g*, along with a part of temporal profiles of the seven cells numbered 1-7. (i-j) Intracellular propagation of [Ca^2+^] observed at 9.4 fps. (i) A raw image in the region indicated with a dashed light-blue square in (a), with the closeups just before and after the [Ca^2+^] increase. (j) Emergence and propagation of [Ca^2+^] within a cell displayed with an intensity relative to the baselines of temporal profiles at each pixel. (k) Temporal profiles with offsets at five positions indicated with *I*-*N* in the left panel of (j).

The randomness of the spatiotemporal distribution of pulsing cells was evaluated by the nearest-neighbor (NN) analysis,^23^ as shown in Fig. 4g-bottom. It clarified that pulsing occurred randomly in most of time and place, but clustering of pulsing cells was found at three local spatiotemporal regions, *F*, *G*, and *H*, shown in Fig. 4f-g. Videos of these specific regions revealed that the clustering was caused by [Ca^2+^] propagation from a cell to surrounding cells (Video 2, panel *iii* and *iv*). Four frames of the [Ca^2+^] propagation at the location *G* are shown in Fig. 4h, along with a part of temporal profiles of seven cells numbered 1–7. The time profiles clearly show how the pulses propagate in sequence from the central cell to the distal cells. Almost the same phenomenon was observed upon addition of high histamine (Video 1). Common to these phenomena, the wave propagation appeared to be triggered by the loss or shrinkage of a cell near the center. One possible explanation for this phenomenon is that intracellular ATP is released into the extracellular space from stressed cells such as mechanically-stimulated cells^24^ and necrotic cells^25^ and binds to the ATP receptors (P2Y) in surrounding cells, causing an increase in [Ca^2+^]. However, in all cases, the propagation distance was within a 10-cell radius. Since the released ATP diffuses into the environment and decreases distally, this result may suggest that above a radius of the 10 cells, the [ATP] is below the threshold for HeLa cells to respond.

In addition, we demonstrated that the intracellular [Ca^2+^] propagation within a single cell could be observed at a frame rate of 9.4 fps (106.5 ms interval), which was the highest frame rate of the imaging system (Fig. 4i-j). Instead of the ratiometric detection of two-wavelength channels, calculating the ratio of the FRET channel intensity to its baseline in the temporal profiles at each pixel (Fig. 4k) enabled the visualization of Ca^2+^ propagation within 20 frames (∼ 2 s) (Fig. 4j, Video 3). We would like to restate that the observation of intracellular spatiotemporal dynamics was achieved with an over-centimeter FOV.

### Rare-cell-triggered macroscale pattern formation of *D. discoideum*

The ability of AMATERAS1.0 to detect rare and functionally important cells was demonstrated by the observation of a multi-cellular dynamical system. *D. discoideum* cells are known to change their life style from uni-cellular to multi-cellular mode in their development initiated by the environmental stress such as nutrient depletion. In this process, these cells microscopically initiate the intercellular relay of cAMP, being a chemoattractant molecule, and macroscopically self-organize the aggregation stream in the form of spiral and circular waves over a centimeter range. Many studies have been conducted aiming to elucidate the pattern formation mechanism,^26–28^ but a consensus has not been established with respect to several hypotheses.^29–31^ One of crucial subjects is which cell and by what mechanism triggers the behavior of other cells in the early stages of development of macro-scale patterns. Here, we demonstrate that AMATERAS1.0 can contribute to this subject, as it can be used for simultaneous imaging of single-cell signaling and global pattern formation, in contrast to previous studies where the imaging was independently employed either with a wide FOV and low-spatial resolution or that with a narrow FOV and a high-spatial resolution.

To visualize and quantify [cAMP], we specifically designed and constructed *D. discoideum* cells expressing a ratiometric fluorescent indicator, a fusion protein Red-FL2 of Flamindo2^32^ and a monomeric red fluorescent protein (mRFP),^33^ which are sensitive and insensitive to [cAMP], respectively. Dual-color mapping allows for the visualization of low-[cAMP] cells in yellow color and high-[cAMP] cells in red color. We cultured clonal population of the above cell line, and replaced the culture medium by a development buffer at a time point (*t* = 0) to set a starvation condition to trigger the pattern formation. Figure 5a shows the time evolution of the macroscale pattern of cAMP signaling at five time points in more than 15 hours of the measurement period. The number of cells observed in the FOV was approximately 2.4 × 10^5^ at the beginning of the measurement (*t* ∼ 0). We were able to observe the transition of the macroscale pattern starting from the non-wave state (Fig. 5a, left and 2nd-left) followed by a spiral wave (center) and a circular wave (2nd-right). Finally, the cells migrated toward the centers of the circles to form multicellular aggregates (right). In all phases of the macroscale pattern formation, the cAMP of individual cells as well as their movement were observed at sub-cell resolutions (Fig. 5b-d). The spatial resolution (< 2.5 µm) and image acquisition frequency (every 30 s) were high enough to recognize single cell motion and morphology (Fig. S6). An overall video of the full FOV and magnified views can be found in the supplementary materials (Video 4).

**Fig. 5.**
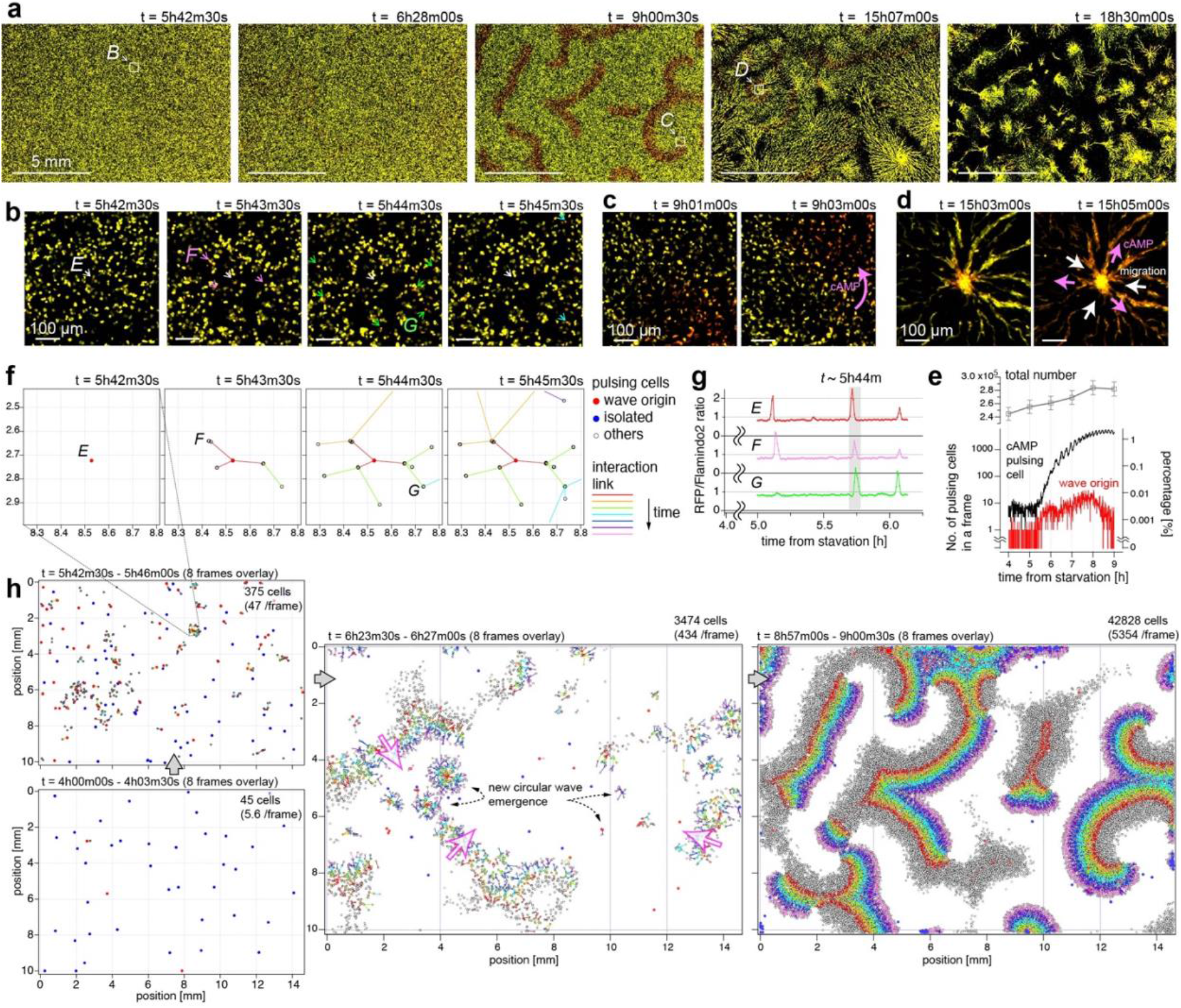
Formation of the spatial pattern of *D. discoideum* on a centimeter scale. The *D. discoideum* cells express two fluorescent proteins, namely, Flamindo2 and mRFP, which are sensitive and insensitive to [cAMP], respectively. Images were recorded every 30 seconds over 16 hours after 4 hours following cellular starvation (*t* = 0h00m). (a) Evolution of macroscale pattern on a large time scale. The images are shown in 8bit-RGB, in which the green and red colors reflect the intensity of Flamindo2 and mRFP, respectively. The color scales were adjusted so that the cell color changed from yellow to red upon an increase in the [cAMP] from low to high. The time counters indicate the time conceded since the starvation. (b-d) Closeups of three locations indicated by dashed squares in (a) at the three time-regions; region *B* at *t* ∼ 5h44m (b), *C* at *t* ∼ 9h01m (c) and *D* at *t* ∼ 15h03m (d). (b) The central cell, denoted with a white arrow, originated a [cAMP] wave, which propagated to several cells denoted with pink, green, and light-blue arrows. (c-d) Propagation of [cAMP] and cell migration are indicated with pink and white arrows, respectively. (e) Time evolution of the number of pulsing cells (black), cells working as wave origins (red), all cells (gray). Their percentage in total cells are represented by the right axis. (f) Distribution of auto-detected cell positions with tree-network diagram of [cAMP] propagation at the same location and frames as (b). (g) Temporal profiles of the fluorescence intensity ratio (mRFP/Flamindo2) of the three cells marked with *E*-*G* in (f). (h) Scatter plots of cell positions across 8 successive frames with their tree-network diagram in the full FOV at four time-regions.

We now focus on the early time region until *t* = 9h00m in which a macroscale spiral wave was stably generated from the non-wave state. By performing an image analysis of the image sequence observed in the full FOV, we detected all pulsing cells among more than 2.4 × 10^5^ cells. Figure 5e shows the time evolution of the number of pulsing cells (black), where those of total number of all cells (gray) and the number of cells working as wave origins are also shown. Before *t* = 5h30m, the number of pulsing cells in one frame was less than 10 (< 0.004 % of 2.3 × 10^5^ cells) in most frames (Fig. 5e). After *t* = 5h30m, the number of pulsing cells began to increase, and the wave propagation began to appear. As an example, around *t* = 5h42–45m, while no macroscale wave was observed yet in the full FOV image, one can see in a local region *B* (Fig. 5a left) that a pulsing of the central cell was followed by several surrounding cells with a delay of a few frames (Fig. 5b). This observation can be attributed to the propagation of the wave owing to the triggering of the surrounding cells by the central cell. In order to visualize the causal relationship of the pulsing cells more clearly, tree-network diagram was created based on the pulsing order of neighboring cells. The tree-network for the same area of Fig. 5b is shown in Fig. 5f. The temporal profile of [cAMP] in the three cells (*E*-*G*) also indicates that the central cell (*E*) is the origin of the wave in this local region (Fig. 5g). The tree-network diagrams in the entire FOV at four time points are shown in Fig. 5h, including the spatiotemporal region of Figs. 5b and 5f (See also Video 5). Deciphering such network diagrams can lead to an understanding of the mechanism of macro-scale wave generation. For example, it is possible to identify the cells that are the origins of waves, as shown in Fig. 5e.

In the image sequence, we also found another rare phenomenon that a very small fraction of the cells cannibalized their neighboring cells. This phenomenon, called entosis, was first studied on epithelial cells detached from extra-cellular matrix^34^ and recently its function in cancer has been well studied.^35^ Cannibalization of *D. discoideum* has been discussed for much longer than in other cells, but it was understood as a process when a giant cell engulfs hundreds of neighboring cells to form a macrocyst.^36,37^ In the current study, we observed entotic phenomena occurring in the starved condition which seems different from the one leading to the macrocyst formation.

Figure 6a shows an image sequence involving an entotic event, which was found at *t* = 5h50m, where a cell marked with *A* came into contact with the other cell *B* and eventually engulfed it into the body. It was noticed that the entotic cells exhibited two features in the appearance; The color inside of the combined cell turns red, as can be seen in Fig. 6a and other examples in Fig. 6b, and the red color typically lasts much longer than the time duration of the cAMP pulse (∼ 2 minutes, Fig. 5g). Figure 6c shows the histogram of time duration of the “red-cell”-encapsulated state, where we found that the distribution of time duration decreases monotonically like an exponential distribution and some cells have time duration even longer than 1 hour. The color appearance is caused by the lower fluorescence intensity of Flamindo2 (green channel of the dual-color image) presumably due to the lower pH (∼ 5) inside the phagosome^38^ generated during the entotic process. Since the fluorescence intensity of Flamindo2 is pH-sensitive (p*K*a > 7.5),^32^ whereas mRFP has a much less dependence on pH (p*K*a = 4.5),^33^ the mRFP/Flamindo2 ratio increases, resulting in the appearance of red particles inside.

**Fig. 6.**
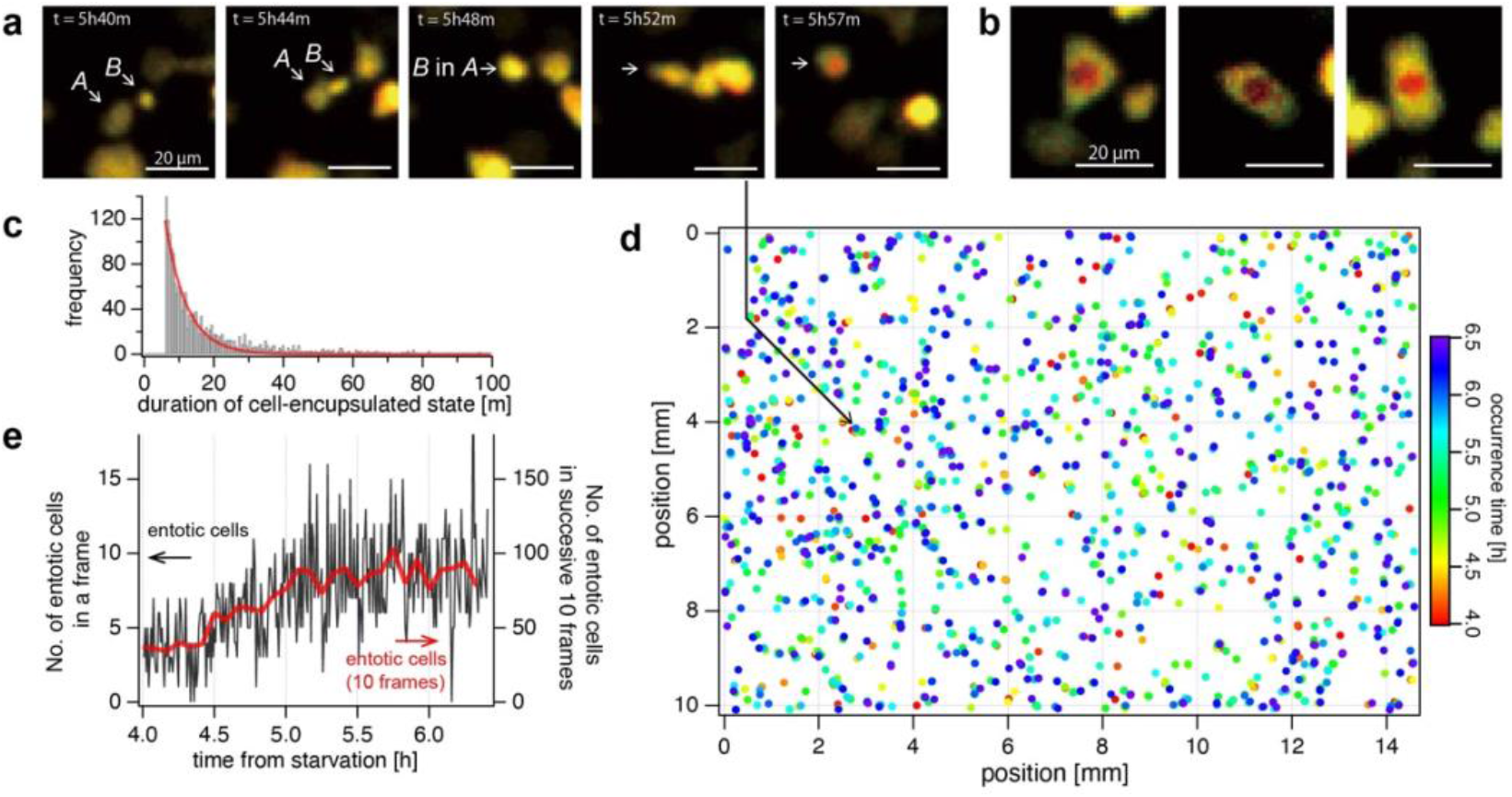
Detection and analysis of entotic event of *D. discoideum*. Entotic events were detected in the same image-set as Fig. 5 from 1st to 300th frame (*t* = 4h00m – 6h30m). (a) An example of entotic event observed from 5h40m – 5h57m. The larger cell marked with *A* came into contact with the smaller cell marked with *B*, and the two cells have unified at *t* = 5h48m. Within the next ten minutes, the inside of the unified cell turned red. (b) Three examples of the typical color appearance of entotic cells. (c) Histogram of the time duration of “red-cell”-encapsulated state. (d) Spatiotemporal distribution of the occurrence of detected entotic events in the entire FOV. The Rainbow color table represents the time of occurrence. (e) The temporal evolution of the number of entotic events detected in each frame (black line) and in successive 10 frames (red line).

We created a spatiotemporal map of the occurrence of this phenomenon, as shown in Fig. 6d and Video 6. The number of the entotic events occurring in each frame is plotted in Fig. 6e. The percentage of the entotic cells in all cells was found to be 1.4 % over the 2.5 hours, which corresponds to 0.0048 % per frame. The spatiotemporal distribution of the occurrence of the entotic event appeared to be random, but its frequency gradually increased as time progressed even before the number of cAMP pulsing cells started to increase (Fig. 5g). The increase of the entosis frequency may be attributed to their nutritional intake to survive the starvation, which was suggested in the previous studies on other types of *Dictyostelium*.^39,40^

## Discussion

In this paper, we established an imaging system AMATERAS1.0 that allows spatially and temporally resolved observation of single cells within the over-one-centimeter FOV, and showed its possibility to detect and analyze rare and unique cells among numerous cells that are observed simultaneously. In the imaging of MDCK cells (Fig. 3), we have demonstrated that more than one million cells can be measured in a single-shot imaging with one-second exposure time. The number of observable cells depends on cellular size and density, of course. In eukaryotic cells, the maximum number of cells is thought to be about 1 million, as shown in this study. However, since prokaryotic cells are smaller in size, more than 10 million cells are thought to be possible in principle. By measuring this large number of cells (10^6^ or more), 0.0001% of rare cells can be detected. If the rarity is 0.01%, it is possible to detect about 100 of these cells, and statistical analysis can be performed to understand their behavior and functions.

Furthermore, we verified the ability of AMATERAS1.0 for high-throughput single cell analysis. Specifically, as a proof-of-concept, we performed imaging of nucleus-stained MDCK cells to study both the populations and spatial distribution of cellular state, *i.e.,* cell cycle phases, of individual cells. Indeed, it was found that cells in the same phase have a common tendency to cluster with each other at the two different cell densities, and furthermore, the spatial scale of the clusters were found to be different between the two conditions. The above result indicated the promise of AMATERAS1.0 as a high-throughput cytometry method. We would like to emphasize that it would be a powerful tool for the statistical analysis of cellular states in the study of the cellular heterogeneity and diversity. ^41,42^ Flow cytometry, which is widely used for cellular statistics, has a throughput of 10^2^-10^3^ cells per second. A throughput of >10^4^ cells per second has been realized with other state-of-the-art technologies.^43,44^ In contrast, our imaging system has the potential to be the next-generation tool for imaging cytometry with a throughput of >10^5^ cells in a single shot. Moreover, AMATERAS1.0 can perform time-lapse cytometry for dynamic transition of cellular state in a multi-cellular system,^45^ which is impossible by means of the flow cytometry. As a possibility for further development, combining MERFISH (multiplexed error-robust fluorescence hybridization), which is known as a transcriptome method under imaging, ^46^ is promising to maximize the advantage of image cytometry.

In the imaging of spontaneous [Ca^2+^] elevation in HeLa cells (Fig. 4), we have shown that AMATERAS1.0 can detect spontaneous pulsing that occurs only in much less than 1 % of rare cells. This study has quantitatively evaluated the frequency (rarity) of the spontaneous increase in [Ca^2+^] for the first time. It has recently been suggested that spontaneous increases in [Ca^2+^] may contribute to the regulation of progenitor cell growth in cardiac and neuronal tissue.^47,48^ However, the biological role of the spontaneous [Ca^2+^] pulsing in HeLa cells, which are non-excitable cells unlike the cardiac or neuronal cells, has not been studied. A high-throughput imaging system such as our system would be indispensable to study the significance of such rare events together with their mechanisms.

In the *D. discoideum* imaging, we focused on [cAMP] elevation (Fig. 5). We detected all cells with elevated [cAMP] during the time period from starvation (*t* ∼ 0) to the phase when the spiral wave is stably repeated (*t* ∼ 10h), and clarified the changes in their frequency and spatial distribution. Since the purpose of this paper is to demonstrate the methodology, we only present the results of measurement and analysis. In another paper submitted elsewhere, we analyzed the same kind of dataset in detail and compared it with theoretical simulations to elucidate the mechanism of spiral wave generation.^49^ Briefly, we found that the non-uniform distribution of pulsing cells at the time region around *t* ∼ 6h, which is incidental due to their rarity (< 0.02 % in a frame), is a key factor for the spiral wave formation at the time region around *t* ∼ 9h.

We have also demonstrated the detection of entotic cells, and clarified their frequency (Fig. 6). Although the role of entosis in *D. discoideum* has not been well known, the present results have provided the first quantitative analysis of the rarity of the spatiotemporal distribution of entotic phenomena in *D. discoideum*. In the future, we would like to elucidate their biological significance by adding other feature parameter in the imaging.

We believe that by observing such a large number (10^5^-10^6^) of cells in live imaging, we will be able to unravel the existing mysteries about rare cells and also discover new research targets. In order to promote this, the instrument needs to be widely available. As mentioned in the introduction, AMATERAS1.0 is a simple system mainly consisting of a low-cost lens and camera available on the market. This feature ensures that it can be easily implemented in small laboratories and is not too difficult for general biologists to build. Customization of the system do not require a high level of optical expertise, so that it can be modified according to the research targets of individual researchers. Therefore, effective dissemination of this powerful tool in biological laboratories can be achieved, which would contribute to the creation of a new scientific field.

## Methods

### Imaging system: AMATERAS1.0

The imaging system (AMATERAS1.0, Fig. 1a) was built using the DIY components provided by Thorlabs and OptSigma. We used a vertical support rail (CEA1600, Thorlabs, Newton, NJ) as the main body. An inverted microscope configuration was employed for imaging of the cultured cells. A camera (VCC-120CXP1M, CIS, Tokyo, Japan) equipped with the 120-megapixel image-sensor (120MXSM, Canon, Japan) was used. A 2× macro-lens was used as imaging lens for a full-size sensor (LSTL20H-F, Myutron, Tokyo, Japan). Owing to its long working distance, the sample was illuminated obliquely with fluorescent excitation light from the bottom, which allows for epi-fluorescence imaging without a dichroic mirror. The incident angle was set to 35°, which was higher than the angular aperture of the imaging lens (∼ 7°) to prevent direct reflection from entering the imaging lens. For fluorescence excitation, we used four high-power LEDs with center wavelengths of 525 nm, 470 nm, 445 nm, and 385 nm (SOLIS-525C, SOLIS-470C, SOLIS-445C, and SOLIS-385C, Thorlabs, Newton, NJ) along with corresponding four sets of excitation filters (#86-963, #86-962, #86-961, #33-322, Edmund Optics, Barrington, NJ) and fluorescence filters (#67-048, #86-366, #67-042, #84-111, Edmund Optics, Barrington, NJ) with a 2-inch diameter. The excitation filters were placed right after the LED sources. The LED emissions were combined at two dichroic beam-combiners (#86-399, #86-396, Edmund Optics, Barrington, NJ) along the same path. The light is relayed by a lens system composed of two lenses with 0.5X magnification, namely, L1 (f = 150 mm, LA1417-A-ML, Thorlabs, Newton, NJ) and L2 (f = 75 mm, LA1145-A-ML, Thorlabs, Newton, NJ), and reflected by a mirror toward the bottom surface of the sample. The LEDs were switched on/off by a trigger signal from a computer through a DAQ board (NI USB-6001, National Instruments, Austin, Texas). The fluorescence filters were switched by a motorized wheel used for a 2-inch filter (#59-769, Edmund Optics, Barrington, NJ). However, the two fluorescence filters (#67-048, #86-366) were switched to a filter flip mount (MFF102/M, Thorlabs, Newton, NJ) for high-speed switching only for the imaging of *D. discoideum* cells. To control the focus position, the lens and camera were mounted on a single translation stage (PT1/M, Thorlabs, Newton, NJ) with a motorized actuator (SOM-C25E, OptoSigma, Tokyo, Japan), and the entire stage was moved up and down to adjust the focus position. A stage-top incubator (U-140A, BLAST, Kawasaki, Japan) was used for the sample stage to control the temperature and CO2 concentration. The position and angle of the sample stage were adjusted by a five-axis manual stage composed of three translation stages (TSD-651C25-M6, TSD-651C-M6, TASB-653-M6, OptoSigma, Tokyo, Japan) and two goniometer stages (GOHT-65A50BMS-M6 and GOHT-65A75BMSR-M6, OptoSigma, Tokyo, Japan). In particular, fine angle adjustment by the goniometer stages was indispensable for imaging in the wide FOV. The angle was adjusted so that the height difference was within the depth of focus at each end of the FOV. The stage, lens, and camera were housed in a dark box to block out the background light and to suppress the influence of fluctuations in the ambient temperature.

### Preparation of mouse brain slice

Male C57BL/6J mice that were 7-weeks-old were purchased from SLC (Shizuoka, Japan) and used for experiments at least 1 week after animal transportation. Mice were maintained in group housing (usually *n* = 3–6 per cage) with a 12-h light-dark cycle (lights on at 8:00 a.m.) in controlled room temperature. Water and food (CMF, Oriental Yeast, Osaka, Japan) were available ad libitum. All animal care and handling procedures of mice were approved by the Animal Care and Use Committee of Osaka University. All efforts were made to minimize the number of animals used.

For the viral construct, pAAV-mDlx-GFP-Fishell-1 was kindly provided by Gordon Fishell (Addgene plasmid #83900; http://n2t.net/addgene:83900; RRID: Addgene_83900). The AAV plasmid vector including the mouse alpha-CaMKII promoter was kindly provided by Akihiro Yamanaka (Nagoya University), and pAAV-CaMKII-tdTomato-WPRE was constructed by the insertion of the tdTomato open reading frame.

AAV packaging and titration were performed as previously described^50^ with minor modifications. Briefly, AAV transgenes were packaged using an AAV helper-free packaging system (catalogue no. VPK-400-DJ; Cell Biolabs, San Diego, CA, USA) except for the plasmid carrying AAV rep and cap genes. A plasmid vector carrying the AAV PHP.eB capsid was synthesized by Ken-ichi Inoue (Kyoto University) and Masahiko Takada (Kyoto University) according to a previously published report outlining the method for the development of PHP.eB capsid. The AAV transgene plasmid, AAV helper plasmid, and AAV rep and cap plasmid for the construction PHP.eB capsid were co-transfected into HEK293T cells using polyethyleneimine (catalogue no. 24765; Polyscience, Inc., Warrington, PA). The cells and culture media were separately harvested 72 h after transfection, and crude AAV preparation was obtained after the cell suspension was subjected to the freeze-thaw cycle in lysis buffer (10 mM Tris, 10 mM MgCl2, and 150 mM NaCl, pH 7.6) and polyethylene glycol precipitation from the culture media. Crude AAV preparation was treated with ≥250 units/μL Benzonase® Nuclease (catalogue no. E1014; Sigma-Aldrich, St Louis, MO), and the supernatant was collected after centrifugation at 3,000 × g for 15 min at room temperature. Viral vectors were purified using iodixanol (Optiprep, catalogue no. AXS-1114542; Cosmo Bio Co., Tokyo, Japan) density gradient ultracentrifugation.

The viral titers were determined by quantitative real-time PCR using GoTaq® qPCR Master Mix (Promega, Madison, WI) on a CFX96 Touch™ Real-Time PCR Detection System (Bio-Rad, Hercules, CA) with a linearized pAAV-mDlx-GFP-Fishell-1 as a standard.

### Viral injection and tissue preparation

Mice were intravenously injected with a virus cocktail containing 1×10^11^ genomes of viruses AAV-PHP.eB-CaMKII-tdTomato-WPRE and AAV-PHPeB-mDLX-GFP. Three weeks after viral injection, mice were deeply anaesthetized by intraperitoneal injection of an anaesthetic cocktail comprising 4 mg/kg midazolam (Dormicum, Astellas, Tokyo, Japan), 0.3 mg/kg medetomidine (Domitor, ZENOAQ, Fukushima, Japan), and 5 mg/kg butorphanol tartrate (Vetorphale, Meiji Seika Pharma, Tokyo, Japan), and were transcardially perfused with 10 ml saline followed by 15 ml of 4% paraformaldehyde dissolved in phosphate-buffered saline (PBS). Brain tissues were excised and stored in 4% paraformaldehyde dissolved in PBS at 4 °C overnight and transferred to 0.05% solution of sodium azide dissolved in PBS.

Tissue sections were prepared with 25-μm-thickness using a vibrating microslicer (LinearSlicer Pro7N, Dosaka EM, Kyoto, Japan). After nuclear staining with 1 μg/mL Hoechst 33258 dissolved in PBS, tissue sections were mounted on glass slides and coverslipped with ProLong Glass antifade mountant (catalogue no. P36982; Thermo Fisher, Waltham, MA).

### Preparation and imaging for MDCK cells

The MDCK cells were cultured in a 35 mm glass-bottom dish (AGC Techno Glass, Shizuoka, Japan) up to the interval immediately before reaching confluence (pre-confluent) or 1.5 day after becoming confluent (post-confluent) at 37 °C with 5% CO_2_ in D-MEM (043-30085, Wako, Osaka, Japan) supplemented with 10% FBS (FB-1365, Biosera, France), 100 units/ml penicillin and 100 µg/ml streptomycin (168-23191, Wako, Osaka, Japan). For nuclei observation, cells were fixed with 4% paraformaldehyde for 10 min, washed with PBS, permeabilized with 0.2% Triton X-100 for 10 min, washed again with PBS, and then stained with 5 µM NucleoSeeing (Funakoshi, Tokyo, Japan) in PBS. Images were obtained using the AMATERAS1.0 system with an excitation LED wavelength of 470 nm.

### Image analysis of the MDCK cells

For images of nucleus-stained MDCK cells, nuclear regions were extracted and background regions were removed using the Pixel Classifier function of machine learning software AIVIA 9.0 (DRVision Technologies LLC, Bellevue, WA). Next, the Nuclei Count function was applied to the images of nuclei to calculate the total fluorescence intensity, centroid position, and size of individual nuclear regions. To visually display the cell state (cell cycle phase) in the right panels of Fig. 3d-e, color maps were made in which every pixel in the individual cell nuclei was filled with a solid 8-bit number corresponding to the total fluorescence intensity of that nucleus.

Spatial distribution of the cells was analyzed in the following two ways, (1) in-phase cell rate of neighboring cells by order of distance and (2) in-phase cell rate in neighboring area as a function of radius of the area (Fig. S4). For the first method, we sampled a cell and sorted the neighboring cells in order of distance from the sample cell, and examined whether they were in phase with the sample cell. The examination was performed for arbitrarily sampled 1000 cells from each of the two cell groups (*G1*, *G2* phases), and the percentage of the in-phase cell is plotted in Fig. 3f. For the second method, we arbitrarily sampled a cell and calculated the rate of in-phase cells within a neighboring region with a radius from the sample cell. The radius dependence of this in-phase cell rate was calculated for 1000 arbitrarily sampled cells from each of the two cell groups (*G1*, *G2* phases), and plotted in Fig. 3g. Here, in both of the analyses, the cells were categorized into the two groups (*G1*, *G2*) based on the midpoint value of the two peaks, although the cells in *S*-phase and *M*-phase are involved in either groups.

### Preparation and calcium ion imaging of HeLa cells

The HeLa cells stably expressing YC3.60 cells were cultured in a 35 mm glass-bottom dish (AGC Techno Glass, Shizuoka, Japan) coated with Cellmatrix Type I-C (Nitta Gelatin, Osaka, Japan) at 37 °C with 5% CO_2_ in FluoroBrite DMEM (A1896701, Thermo Fisher Scientific, Massachusetts, USA) supplemented with 10% FBS (FB-1365, Biosera, France), 4 mM GlutaMax (35050061, Thermo Fisher Scientific, Massachusetts, USA), 100 units/ml penicillin, and 100 µg/ml streptomycin (168-23191, Wako, Osaka, Japan) until the cells reached the confluence. Before imaging, the medium was replaced with FluoroBrite DMEM without FBS. Images were obtained using the AMATERAS1.0 system with an excitation LED wavelength of 445 nm (SOLIS-445C, Thorlabs, Newton, NJ) and an emission filter with a center wavelength of 540 nm (#86-366, Edmund Optics, Barrington, NJ) for imaging in the FRET channel. Two hundred frames were acquired at 5 s intervals (0.2 fps) with 500 ms exposure and at intervals of 106 ms (9.4 fps). The dish was stored in a stage-top incubator at 37 °C with 5% CO_2_. The intensity of the excitation light (LED, 445 nm) in the sample plane was 25.8 mW/cm^2^ and 40.8 mW/cm^2^ at 0.2 fps and 9.4 fps, respectively.

### Analysis of the calcium imaging data

To search for rare cells from 1.2 × 10^5^ cells that spontaneously generate Ca^2+^ pulses, the following processing was performed with a series of images. For all pixels in the FOV, temporal profiles were created and their baselines were flattened with polynomial fitting. A map of skewness of the baseline-corrected temporal profiles was prepared, and particles with large positive skewness were considered as candidate cells. For candidate time profiles, the times of the positive steep peaks were recorded above the noise level along with the coordinates. The markers were then plotted on the images in the full FOV, as shown in Fig. 4f, in which the markers were colored using a red-white-blue color-table based on the time of pulsing. The number of pulsing cells is shown in a histogram according to the peak time (Fig. 4g, top), where the time bin of the histogram was set to 30 s (6 frames).

We introduced the nearest-neighbor (NN) analysis to evaluate the non-randomness of the spatial distribution of pulsing cells (Fig. 4g, bottom). In each frame, the average distance of the pulsing cells from their nearest neighbor (ANN), *W*^(*i*)^, was calculated using the following equation:

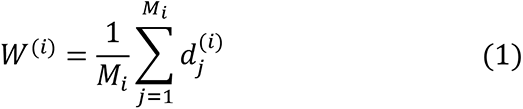

where *M_i_* and *d_j_*^(*i*)^ denote the number of pulsing cells in the *i*-th frame and the distance from the *j*-th cell to its nearest neighbor. Since the number of pulsing cells varies with time, ANN is normalized by dividing with the expected value for obtaining random distribution.

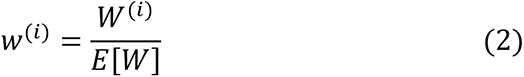

with

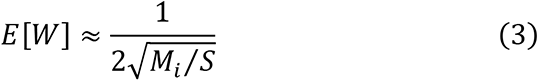

where *S* denotes the area of the FOV, which is constant for all the frames (= 14.6 × 10.1 mm^2^).

When the points are randomly distributed, the ANN (*W*) is known to conform to the following normal distribution, where the *L* and *n* denote the perimeter and the total number of cells.

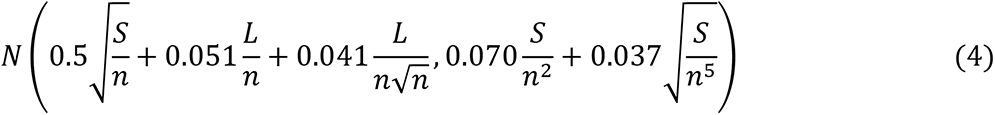

We introduced information content as a measure of non-randomness, given by

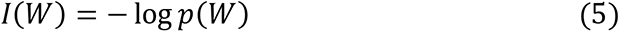

where *p*(*W*) represents the cumulative probability from negative infinity to *W* under the normal distribution obtained by Eq. 4.

In order to visualize the temporal variation of intracellular Ca^2+^ distribution, we obtained a ratiometric image display (Fig. 4j). Since this study uses a single wavelength channel, it is not possible to display the image with the ratio of the two-wavelength channels. Instead, we reconstructed the intensity distribution image by transforming the temporal profile at each pixel (*e.g.*, those in Fig. 4k) in the cell region into its ratio to the respective baseline, for example, 0–4 s in Fig. 4k.

### Preparation and imaging of *D. discoideum*

Ax2 cells expressing the ratiometric fluorescent indicator (fusion protein of Flamindo2 and mRFP) were maintained in a 90 mm plastic dish at 22 °C in HL5 medium supplemented with 50 µg/mL hygromycin (Wako, Osaka, Japan) and 16 µg/mL G418 (Wako, Osaka, Japan). Cultured cells were suspended by pipetting on the dish and split into a new dish at an approximate ratio of 1:6 every 24 h, and the cells were maintained at an almost constant cell number for a long time. Development of the cells was initiated at t = 0h by inducing starvation after washing gently thrice with the development buffer (5 mM Na_2_HPO_4_, 5 mM KH2PO4, 1 mM CaCl_2_, 2 mM MgCl_2_, pH 6.4) to avoid detaching from the dish surface. These cells were suspended in the developmental buffer and the cell number was counted using Countess-II (Thermo Fisher Scientific, Massachusetts, USA). Cells (1.8 × 10^6^) were plated in a 35 mm glass-bottom dish (AGC Techno Glass, Shizuoka, Japan) in 1.6 ml of developmental buffer.

In the fluorescence imaging, two LED light sources with center wavelengths of 470 nm and 525 nm (SOLIS-470C and SOLIS-525C, Thorlabs, Newton, NJ) were used for the excitation of Flamindo2 and mRFP, respectively. The sample dish was placed in a stage-top chamber at 22 °C and imaged by the AMATERAS1.0 system for 16 h from *t* = 4h to *t* = 20h. The imaging interval of the image sequence was 30 s, resulting in 1,920 frames for 16 hours observation. At each time frame, the fluorescence of Flamindo2 and mRFP were sequentially imaged by switching the LEDs and emission filters. The excitation intensities of the two LEDs were 13.4 mW/cm^2^ (470 nm) and 15.4 mW/cm^2^ (525 nm) in the sample plane, respectively. Both fluorescence signals were detected with a 1.3 sec exposure with 8× gain.

### Analysis of the *D. discoideum* imaging data

The total number of cells in the entire FOV was counted manually as it was found rather better than automatic counting by use of a machine learning software because of difficulty in the segmentation of each *D. discoideum* cell. We employed so-called quadrat method. Instead of actually counting all the cells in the image, we counted the cells in a certain compartment of a small area (quadrat). Assuming that the number of cells was uniformly distributed in the area, the cell number density was calculated from the number of cells in the compartment and the total number of cells in the FOV area was estimated. We performed the cell counting at six time points, *t* = 4, 5, 6, 7, 8, and 9, as shown in Fig. 5e.

For Fig. 5e-f, h, pulsing cells were automatically detected out of the 2.4 × 10^5^ cells, and a tree network diagram was created based on their spatiotemporal relationships. The method is outlined as follows: For the detection of pulsing cells, we first obtained images of the intensity ratio of the mRFP to Flamindo2 so that a cell with high [cAMP] has a high value for the ratio. The ratio was obtained only at the pixels where both intensities were higher than a predetermined threshold. The ratio map was binarized according to a preselected threshold that allows only the higher-value pixels to be extracted. Particle analysis was performed on the binary image to distinguish particles with a size higher than a preselected threshold as candidates for pulsing cells. The above analysis was performed from the 1st frame (*t* = 4h00m) to the 600th frame (*t* = 9h00m) to prepare a list of candidate particles with time and positions. Since the list of candidate particles contains many errors, we screened them based on the following procedure. (1) Cells that are within a possible range of cell migration across the consecutive frames were determined as identical. The maximum cell migration distance in 30 s (between two consecutive frames) was empirically set at 16.5 µm (15 pixels). (2) A candidate particle present only in a single frame was excluded since true pulsing cells should be present across multiple consecutive frames. After the two processes, the list of pulsing cells was completed. The time variation of the number of pulsing cells is shown in Fig. 5e.

To draw the tree-network diagram (Fig. 5f and h), links were observed between cells with [cAMP] propagated across successive frames. Since the actual diffusion of cAMP molecules in the solution cannot be measured, the influence relationship was inferred phenomenologically from the spatiotemporal location of the pulsing cells. When a cell was pulsed, if there were pulsing cells within the spatiotemporal range of possible propagation, the earlier and later cells were determined to be the “parent” and the “child”, respectively. The spatiotemporal range is expressed by two parameters: the pseudo-diffusion coefficient (*D’*), which was defined as the distance that the parent cell can influence, *i.e.*, create a child, in the next frame, and the sustainment time (*F_max_*), which was defined as the number of frames in which the influence lasts. For all pulsing cells, we searched for cells pulsing in the subsequent Fmax frames within the diffusion distance derived by *F*^1/2^*D’* (*F*: 1 … *F_max_*) and determined them to be children. We empirically set the two parameters as *D’* = 250 µm and *F_max_* = 3 in this study. This allowed us to create the tree network shown in Fig. 5f and h.

For the detection of entotic events (Fig. 6), we used the feature that cells engulfed by larger cells turn red. First, similarly to the above method for the detection of high-[cAMP] cells, the intensity ratio images of the mRFP to Flamindo2 were made for all the time frames. Only red particles with a ratio value exceeding a predetermined threshold were extracted, and out of those, particles that were too small or too large were judged as errors and excluded. Particles within 11 × 11 µm (10 pixels) between consecutive frames were recognized as the same cell. Since we empirically found that the red particles produced by entosis are maintained for a longer period of time compared to the pulsing of cAMP, we finally determined that those that were present for more than 10 consecutive frames were entosis.

## Acknowledgements

We would like to thank Mr. K. Yasuda of Canon Marketing Japan for his assistance in selecting the image sensor. We are also grateful to Dr. T. Matsuda of Osaka University for his assistance in the preparation of HeLa-YC3.60 cells. We would also like to thank Dr. Kenji Osabe for his critical reading of this manuscript. This work was supported by a Grant-in-Aid for Scientific Research on Innovative Areas “Singularity Biology (No. 8007)” (18H05415 to KH, 18H05416 to HH and 18H05408 to TN), the Research Program of “Five-star Alliance” in “NJRC Mater. & Dev.” (KH, TN) and Precursory Research for Embryonic Science and Technology (PRESTO) of the Japan Science and Technology Agency (JST) (TI).

## Author contributions

TN conceived and coordinated the project. TI and TK built the imaging system. TK cultured cellular samples and imaging was performed. KS, AK, and HH prepared a mouse-brain section. TN constructed HeLa cells stably expressing YC3.60. KH constructed *D. discoideum* cells stably expressing Red-FL2. TI and TK performed the data analysis. TI, KF, TMW, and TN designed the optical system. TI and TN wrote the manuscript with input from all authors.

## Data availability

All the data shown in this paper are available from the corresponding authors upon request.

## Notes

### Competing Interest Statement

The authors have declared no competing interest.

### Summary of Updates

Title revised; Section on "Single shot detection of more than million cells" added with Figure 3; Section on "Rare event detection by calcium imaging in HeLa cells" added with Figure 4; Section on "Rare-cell-triggered macroscale pattern formation of D. discoideum" updated with Figure 5-6; Videos updated and added; Many minor changes in main text and supplemental files

