## Supplemental document for "Exploring rare cellular activity in more than one million cells by a trans-scale-scope"

Taro Ichimura,* Taishi Kakizuka, Kazuki Horikawa, Kaoru Seiriki, Atsushi Kasai, Hitoshi Hashimoto, Katsumasa Fujita, Tomonobu M. Watanabe, and Takeharu Nagai*

*Corresponding authors:

Taro Ichimura,

Takeharu Nagai,

### Contents

(1) Online movie files

Video 1: Histamine-induced [$Ca^{2+}$] oscillation in HeLa cells

Video 2: Spontaneous [$Ca^{2+}$] oscillation in HeLa cells

Video 3: Intracellular propagation of calcium concentration in HeLa cells

Video 4: Macroscale pattern formation of *D. discoideum* cells

Video 5: Time evolution of single cell scatter plot with a tree-network diagram

Video 6: Entotic events detected in the macro-scale pattern formation of *D. discoideum* cells

(2) Notes

Note 1: Influence of the presence of a glass plate between the sample plane and lens

Note 2: Validation of the FOV and pixelation of AMATERAS1.0 in comparison with typically used sCMOS camera and microscope objective lens

Note 3: Comparison with conventional biological microscopes and recent technologies for trans-scale imaging

Note 4: Calcium oscillation induced by histamine addition

(3) Supplementary figures

Figure S1: Numerical calculation of point-spread-function in the presence of a glass plate.

Figure S2: Imaging of the fluorescent beads with typically used sCMOS camera and an objective lens for comparison with AMATERAS1.0.

Figure S3: Comparative plots of FOV and lateral resolution of imaging systems composed of conventional microscope objectives and image sensors.

Figure S4: Schematic illustration of the two analyses on similarity of neighboring cells.

Figure S5: Histamine-induced [$Ca^{2+}$] oscillation and spontaneous radial propagation observed in YC3.60-HeLa cells.

Figure S6: An image sequence of a local region to show the dynamics of motion, morphology, and mitosis at the single-cell level

(4) References for supplementary materials

| | |
|---|---|
| **Video 1** | **Histamine-induced [$Ca^{2+}$] oscillation in HeLa cells** |
| | Representation of Fig. S4 with a video. The video plays simultaneously in both the panels at a speed of 10 fps.<br>(Left) Full FOV video.<br>(*i* – *ii*) Magnified view of the areas indicated by the dashed rectangles in the left panel.<br>(*iii* – *iv*) Views magnified further to demonstrate the cells of interest in the areas indicated by the dashed rectangles in *i* – *ii*, respectively. |
| **Video 2** | **Spontaneous [$Ca^{2+}$] oscillation in HeLa cells** |
| | Representation of Fig. 4a-h with a video. The video plays simultaneously in all the panels at a speed of 10 fps. The markers *A-D* refer to the same cells as those in Fig. 4a, b and e.<br>(Left) Full field-of-view (FOV) video. The light-blue circles appear in the region upon cell pulsation at the exact time of pulsation.<br>(*i* – *iii*) Magnified views of the three areas indicated by the dashed rectangles in the left panel.<br>(*iv* – *vi*) Further magnified views to exhibit cells of interest, in the areas indicated by the dashed rectangles in *i* – *iii*, respectively. |
| **Video 3** | **Intracellular propagation of calcium concentration in HeLa cells** |
| | Representation of Fig. 4i-k with a video. The video plays simultaneously in all the panels at a speed of 10 fps. |

| | |
|---|---|
| | (Left) Full FOV video.<br>(*i*) Magnified view of the areas indicated by dashed rectangles in the left panel.<br>(*ii*) The view of the area that was magnified further as indicated by the dashed rectangles in *i*.<br>(*iii*) The same area as (*ii*) displayed with an intensity relative to baselines of temporal profile at each pixel. |
| **Video 4** | **Macroscale pattern formation of *D. discoideum* cells** |
| | Representation of Fig. 5a-d with a video. The video plays simultaneously in all panels at a speed of 30 fps. The time counter shows the time in the *hh*:*mm* format that has elapsed since the starvation treatment was performed.<br>(Top left) Full FOV video to exhibit the macroscale pattern.<br>(*i* – *iii*) Magnified views of the three areas indicated by the solid rectangles in the top-left panel. The areas (*i* – *iii*) are centered at the same locations as the regions *B-D* in Fig. 5a. |
| **Video 5** | **Time evolution of single cell scatter plot with a tree-network diagram** |
| | Representation of Fig. 5h (bottom) and Fig. 5a (top) with a video. The video plays simultaneously in both the panels at a speed of 10 fps. The time counter shows the time in the *hh*:*mm*:ss format that has elapsed since the starvation process was performed.<br>(Top) Full FOV video to exhibit the macroscale pattern.<br>(Bottom) Scatter plot of the cell positions in 8 successive frames shown with their links. The red markers represent the pulsing cells generating new waves. |

| Video 6 | **Entotic events detected in the macro-scale pattern formation of *D. discoideum* cells** |
|---|---|
| | Representation of Fig. 6 with a video. The video plays simultaneously in all the panels at a speed of 30 fps. The time counter shows the time in the *hh*:*mm* format that has elapsed since the starvation treatment was performed.<br>(Top) Full FOV video.<br>(Middle) Magnified view of the square region in Top, where entotic events were detected. An entotic event starting at $t$ = 5:38 is indicated by two color arrows (white: engulfing, pink: engulfed).<br>(Bottom) Scatter plot of entotic events in the full FOV. |

### Note 1: Influence of the presence of a glass plate between the sample plane and the lens.

In the AMATERAS1.0 system, a thick glass plate (emission filter) was placed between the lens and sample. This is not preferable since it causes a spherical aberration leading to the degradation of the imaging performance. To evaluate the risks of the degradation of imaging performance, a numerical calculation of the PSF in the presence of the glass plate was performed.

Numerical calculation

Figure S1a shows the schematic geometry of the calculation model. We assume that the imaging lens is aberration-free in the absence of a glass plate. The PSF can be calculated by integrating scalar plane waves converging to the geometrical focus of the lens, $\boldsymbol{r}$ = (0,0,0).[1]

$$U(\boldsymbol{r}) = C \int_{\boldsymbol{k} \in NA} \exp[-i\boldsymbol{k} \cdot \boldsymbol{r}] \, d\boldsymbol{k} \tag{1}$$

This integration can be expressed in the polar coordinate system as

$$U(\boldsymbol{r}) = C \int_{\boldsymbol{0}}^{\arcsin(\boldsymbol{NA})} d\theta \int_{-\boldsymbol{\pi}}^{\boldsymbol{\pi}} \exp[-ik(x \sin\theta \cos\varphi + y \sin\theta \sin\varphi + z \cos\theta)] \, \theta d\varphi \tag{2}$$

It is known that this integral can be reduced at the focal plane ($z$ = 0) to a form of $J_1(u)/u$, where $J_n(u)$ denotes the Bessel function of the first type as a function of the distance from the optical axis, $u$.

In the presence of a glass plate between the lens and focus, the glass plate causes phase retardation and displacement of the optical ray upon refraction, leading to the shift and broadening of the PSF. The ray displacement in the transverse and axial directions ($\Delta \boldsymbol{r}$, $\Delta z$) is expressed by the following vectors with *Snell's* law based on geometrical optics.

$$\Delta \boldsymbol{r}(\theta_1, \varphi) = \overrightarrow{BC} = L(\tan\theta_1 - \tan\theta_2)(-\cos\varphi\, \boldsymbol{e}_x - \sin\varphi\, \boldsymbol{e}_y) \quad (3)$$

$$\Delta \boldsymbol{z}(\theta_1) = \overrightarrow{OO'} = L\left(1 - \frac{\tan\theta_2}{\tan\theta_1}\right)\boldsymbol{e}_z \quad (4)$$

where $L$, $\theta_1$, and $\theta_2$ denote the thickness of the plate, incident angle to the plate, and refraction angle of the plate, respectively (Fig. S1b). The vectors $\boldsymbol{e}_x$, $\boldsymbol{e}_y$, and $\boldsymbol{e}_z$ are defined as unit vectors in the directions of the $x$, $y$, and $z$ axes. The phase retardation ($\delta$) is derived by calculating the difference between the optical paths AB and AC, as expressed by

$$\delta(\theta_1) = k(nAC - AB) = kL\left(\frac{n}{\cos\theta_2} - \frac{1}{\cos\theta_1}\right) \quad (5)$$

The ray is displaced in the transverse direction (Eq. 3) along with phase retardation (Eq. 5) since the glass plate is incorporated in the integral for the absence of the glass plate (Eq. 1-2). The integral can be rewritten as

$$U(\boldsymbol{r}) = C\int_{\boldsymbol{0}}^{\arcsin(\boldsymbol{NA})} d\theta \int_{-\boldsymbol{\pi}}^{\boldsymbol{\pi}} \exp\{-i[\boldsymbol{k}\cdot(\boldsymbol{r} - \Delta\boldsymbol{r}(\theta,\varphi)) + \delta(\theta)]\}\,\theta d\varphi \quad (\boldsymbol{6})$$

The intensity distribution is given by the square of *the absolute value of* $U(\boldsymbol{r})$ with $C = 1$.

Results and discussion

Figures S1c and S1d-e show the calculation results of PSF on the $z$-axis (c) and the $xy$ plane (d–e) in the absence and presence of the glass plate. The calculation parameters, wavelength, glass thickness, and refractive index were set to 520 nm, 2.17 mm (sum of 2.0 mm of the filter and 0.17 mm of the coverslip), and 1.52, respectively.

The axial profile (Fig. S1c) indicated that the spherical aberration caused a large shift (745 μm) of the PSF in the axial direction. This is consistent with the predicted values obtained by the geometrical-optics-based equation, Eq. 4, namely, 742 μm for the central ray ($\theta_1 = 0$) and 748 μm for the peripheral ray ($\theta_1 = \arcsin(NA)$). In contrast to the large shift, the broadening of the PSF is only 2 μm in the axial direction, which is negligible compared to the large PSF width in the axial direction. Figures S1d-e indicate that the PSF in the transverse plane is slightly broadened by ~1 %. Such a small difference can also be considered negligible in practical applications.

We can conclude that the spherical aberration owing to the presence of the glass plate causes a significant displacement of the PSF in the axial direction, but does not cause significant degradation of the imaging performance in this low-NA imaging system.

However, care must be taken while performing multi-color imaging with multiple filters since the PSF shift in the axial direction is dependent on the thickness of the glass plate. Even if two filters have the same thickness on their catalogue, the actual thickness has a deviation large enough to cause a difference in the PSF displacement. Therefore, when we performed time-lapse multi-color imaging, we measured the difference in the focal plane by using fluorescent beads in advance, and the focal distance was shifted by the computer-controlled actuator at the same time as the filter switch at every cycle. For example, we observed that the two filters used for the *D. discoideum* experiment in the main text demonstrated ~65 μm difference in the focal plane in the axial direction. Based on Eq. 4, the 65 μm focus shift corresponds to a 191 μm difference in the thickness of the two filters.

### Note 2: Validation of the FOV and pixelation of AMATERAS1.0 by comparison with typically used sCMOS camera and microscope objective lens

In order to demonstrate the validity of the lens and camera employed in the AMATERAS1.0, a comparative experiment was conducted with a conventional lens and camera. We replaced the camera and lens with those that are typically used and acquired fluorescence images of fluorescent beads with a diameter of 2 μm.

Figure S2a shows an image obtained using the AMATERAS1.0 system. The left large panel displays the image of the full FOV, and the right two panels show the closeups of two local regions, namely, the center (*A*) and the corner (*B*) of the FOV. As shown in Fig. 1b-e in the main text, fluorescent beads can be observed with almost the same spatial resolution even at the corner.

Figure S2b shows an image of the same sample obtained with the imaging lens replaced by a pair of objective and tube lenses (TL2X-SAP, TTL200-A, Thorlabs, New Jersey). The imaging property of this lens pair is characterised by a magnification of 2x, NA of 0.1, and a field number of 22 mm. Because of the narrower field number compared to that of the imaging lens used in AMATERAS1.0, the effective FOV is restricted to the 11 mm circle (dashed yellow circle). The left panel displays the full FOV, and the right three panels show closeups of the three local regions at the center (*A*), the periphery of the FOV (*B*), and the periphery of the image sensor (*C*). The fluorescent beads could be imaged at the peripheral region of the FOV, but the intensity is significantly reduced. The beads could not be imaged at all at the corner of the image sensor.

Figure S2c shows an image of the same sample obtained with the image sensor replaced by an sCMOS camera (Zyla 4.2, Andor, Belfast, UK). The chip size is 18 mm (diagonal), which is much smaller than the image sensor of AMATERAS1.0 (35 mm). Therefore, the FOV is restricted by the CMOS chip. The pixel size of Zyla 4.2 is 6.5 μm, which is approximately three times larger than that of the image sensor of AMATERAS1.0 (2.2 μm). Because of the 2x magnification, the pixelation size is 3.25 μm, which is larger than the spatial resolution of the imaging lens, ~2.25 μm, as described in the main text. Thus, the spatial resolution of this system is determined by the pixel size rather than the NA of the imaging lens.

Through these comparisons, the use of the combination of the imaging lens and the image sensor in AMATERAS1.0 was validated.

### Note 3: Comparison with conventional biological microscopes and recent technologies for trans-scale imaging.

In this paper, we proposed an imaging system for sub-cellular resolution in over-one-centimeter FOV using a hundred-megapixel image-sensor and camera lens designed for machine vision. Fluorescence imaging of live cells has been demonstrated with a single-cell resolution in a much larger FOV than that of the conventional biological microscopes. The spatial resolutions and FOV corresponding to AMATERAS1.0 system and several conventional imaging systems (combinations of the imaging lens and image sensor) are plotted in Fig. S3 for visual comparison, with the vertical axis representing spatial resolution and the horizontal axis representing FOV size (diagonal). We selected five series of objective lenses from three microscope companies (Olympus, NIKON, Thorlabs), and two widely-used image sensors obtained from Andor. See the figure legend for further details.

The effective spatial resolution ($d$) is defined as the larger spatial resolution of the two types with the limit based on optical diffraction ($d_{\text{lens}}$) and pixelation at the image sensor ($d_{\text{sensor}}$).

$$d = \max(d_{lens}, d_{sensor}) = \max\left(C\frac{\lambda}{NA}, 2a\right) \qquad (7)$$

Here, $\lambda$ and $a$ denote the wavelength of light and pixel size of the image sensor, respectively. The coefficient $C$ depends on the resolution criterion (0.5 for Abbe's criterion, 0.61 for Rayleigh's criterion, and 0.47 for Sparrow's criterion). However, the Abbe criterion ($C$ = 0.5) should be adopted here for effective comparison with the sampling rate by the pixelation. In contrast, the FOV size ($\Phi$) is defined as the smaller FOVs restricted by the lens system ($\Phi_{lens}$) and image sensor ($\Phi_{sensor}$).

$$\Phi = \min(\Phi_{lens}, \Phi_{sensor}) = \min\left(\frac{FN}{M}, \frac{\emptyset}{M}\right) \quad (8),$$

where *FN*, $\varphi$, and *M* denote the field number of the lens system, the physical size of the image sensor, and the magnification of the lens system. To represent the ability to cover the scale hierarchies, we introduce "trans-scalability index", *r*, which was defined by the ratio of one-dimensional FOV (Eq. 8) to the lateral spatial resolution (Eq. 7) derived by:

$$r = {}^{\Phi}/_{d} \quad (9)$$

This index has the same function as the space-bandwidth product proposed in previous literature[2] but offers a more intuitive understanding of the scale hierarchy covered by the optical system. In other words, the index represents the number of effective sampling points at which the full FOV is divided by its diameter. Contour lines of the trans-scalability index are shown in Fig. S3.

To achieve a high trans-scalability index (*r*) in wide-field imaging, a large number of pixels is the minimum requirement. At the same time, the chip size must be within the FOV of the lens system, and as a result, a small pixel size is also required. The maximum value for a given image sensor is achieved when the chip size is less than the FOV of the lens system*, and* the pixel size is greater than half of the optical resolution ($d_{sensor} > d_{lens}$). For low-magnification ($M < 10$) lenses, this condition is largely met. Hence, the markers in the plot of Fig. S3 are aligned along the image sensor-dependent contours ($r_{max} = \Phi_{sensor}/d_{sensor}$). The two types of scientific cameras, namely, Zyla 5.5 and iXon Ultra 888 (Andor, Belfast, UK), have chip sizes of 21.8 mm and 18.8 mm, and pixels of 6.5 μm and 13 μm, respectively. Hence, the maximum values of $r$ are $r_{max}$ = 1677 and $r_{max}$ = 723, respectively. This limit cannot be exceeded with any lens system. In contrast, oversampling ($d_{sensor} < d_{lens}$) and the value of $r$ decreases in the high-magnification region ($M > 10$). Compared to these conventional cases, in the present study, a sensor with an extremely high number of pixels was used, which resulted in an $r$-value of 7487. Since the lens system was selected to obtain comparable spatial resolutions of the lens system and image sensor ($d_{sensor} \sim d_{lens}$), it can be said that this is an ideal configuration that achieves a high $r$ without compromising the optical resolution at the sub-cellular scale.

Imaging systems reported by other groups[3,4,5] are also plotted in Fig. S3 for comparison. The values of spatial resolution and FOV reported in the paper were adopted for the plots, even though the performance is still being improved. Our AMATERAS1.0 system outperforms the other groups with respect to the FOV, as it prioritises the expansion of the lateral FOV over resolution. In contrast, the AMATERAS1.0 system was inferior in terms of spatial resolution owing to the low NA of the imaging lens. The advantage of this system is that this performance can be achieved with simple configuration and low-cost devices, which contributes to the potential for wide applications.

### Note 4: Calcium oscillation induced by histamine addition

In the main text, we have shown that HeLa cells in a culture chamber undergo spontaneous elevation of $Ca^{2+}$concentration ($[Ca^{2+}]$) at an extremely low frequency. Here, we present an observation of $[Ca^{2+}]$ oscillations induced by histamine stimulation, which is a well-studied phenomenon.[6,7,8,9,10]

The same HeLa cell samples used in the experiments in the main text were used. The cells were stimulated with histamine (#081-03551, Wako, Japan) at a concentration of 5 μM and time-lapse imaging was performed at an imaging rate of 5 s intervals. The results obtained are shown in Fig. S5. Figure S5a shows the full FOV, and Fig. S5b shows a partial magnification of the area indicated by the white rectangle. In this region, we observed a large number of cells pulsing periodically. In addition, in the time range of $t$ = 420-490 s, we were able to capture a $[Ca^{2+}]$ wave propagating radially from the center to the surrounding area (Fig. S5c). The temporal profile at the seven locations (indicated by *A*-*G*) at the top left of Fig. S5c is shown in Fig. S5d. It can be observed that some of the cells pulsed for a period of approximately 100 s. In contrast, cell *E* caused only a single increase in $[Ca^{2+}]$ and an increase due to $[Ca^{2+}]$ propagation at approximately $t$ = 420 s. Cell *D* was not even affected by $[Ca^{2+}]$ propagation. Thus, it is evident that there are individual differences in calcium properties, probably owing to the cellular arrangement and state. By using our trans-scale-scope, we can statistically investigate such individual differences in cells.

The results of Fig. S5 is also shown in a video (Video 1).

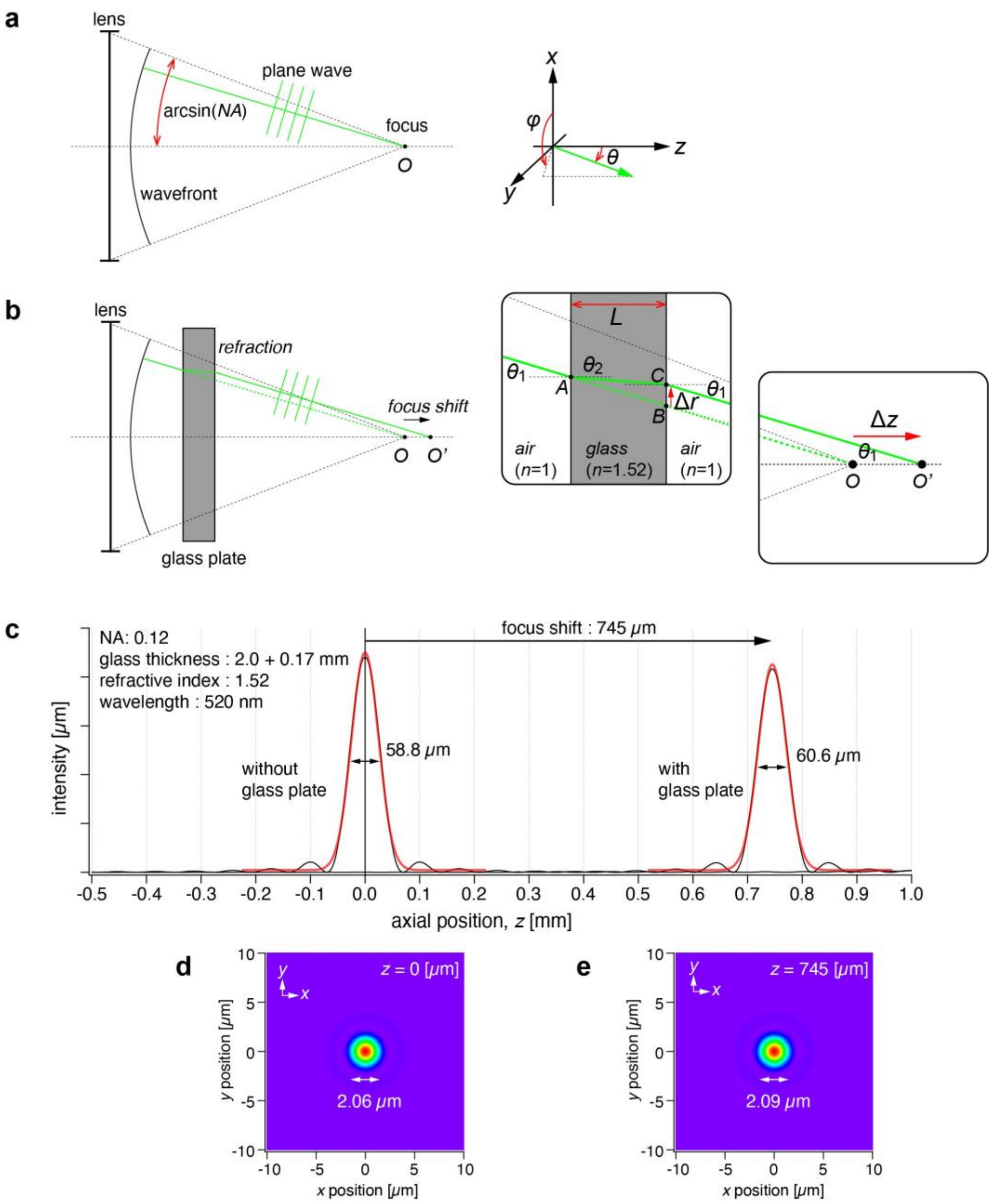


**Figure S1**: **Numerical calculation of point-spread-function in the presence of a glass plate.** (a-b) Schematics of the calculation model in the absence (a) and presence (b) of a glass plate. (c) Intensity profile along the *z*-axis without and with the glass plates, indicating the displacement of the focus position in the z-direction by 745 μm. The wavelength, glass thickness, and refractive index were set to 520 nm, 2.17 mm (sum of 2.0 mm of the filter and 0.17 mm of the coverslip), and 1.52, respectively. (d-e) Intensity distribution in the *xy* plane at the $z$ axial position, (d) $z = 0$ for the case without the glass plate, and (e) $z = 745$ for the case with the glass plate.

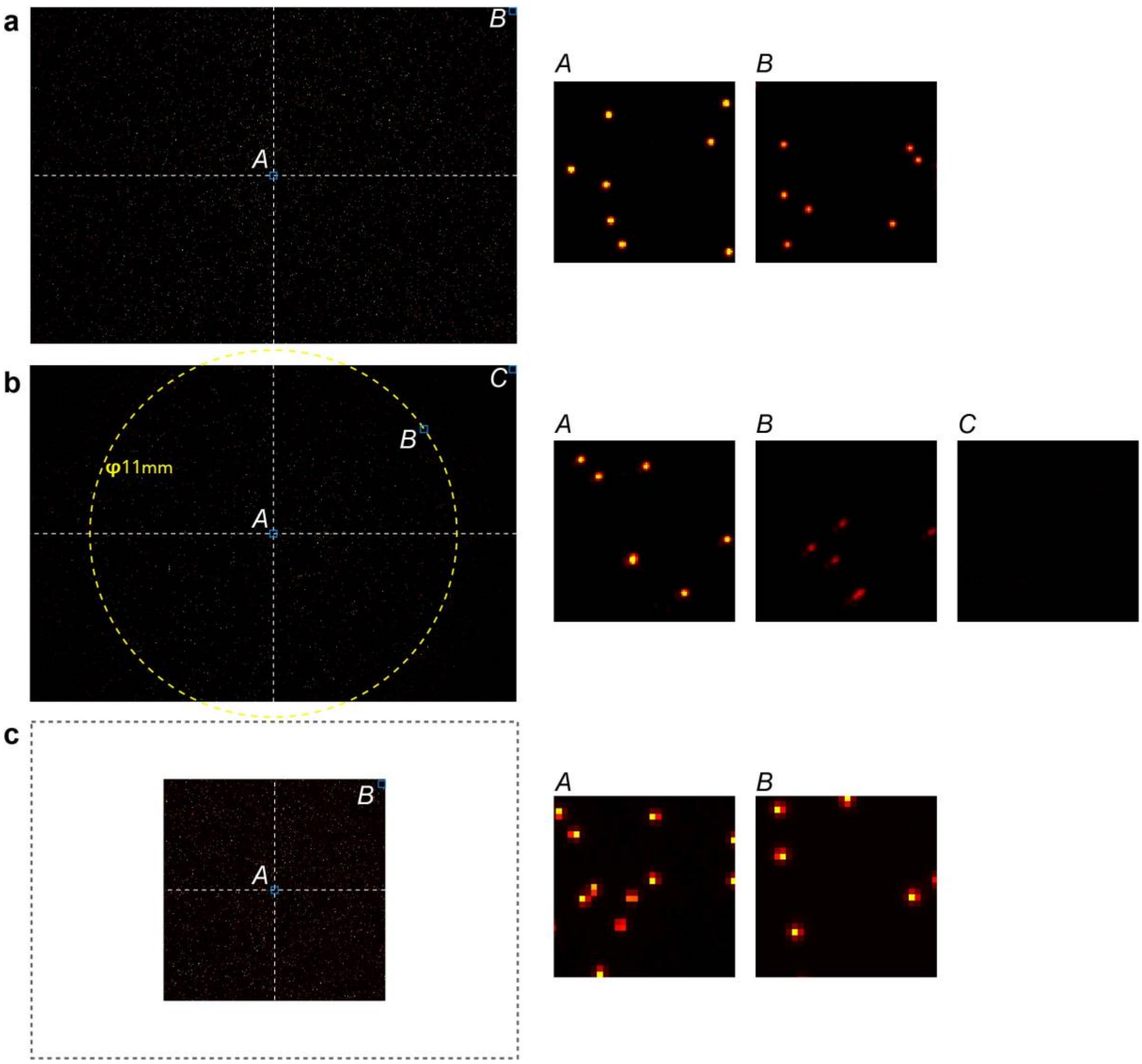


**Figure S2**: **Imaging of the fluorescent beads with typically used sCMOS camera and microscope objective lens for comparison with the AMATERAS1.0 system.** (a) An image obtained by the AMATERAS1.0 system. The left panel shows the full FOV regulated by the image sensor, and the right two panels show the magnified views of the two local regions (*A*: center, *B*: corner). (b) An image obtained by the combination of a 2X imaging system of Thorlabs and the image-sensor used in the AMATERAS1.0 system. The left panel shows the full image obtained by the image sensor, and the right three panels show the magnified views of the three local regions (*A*: center, *B*: an edge of FOV regulated by the lens, *C*: corner). The dashed yellow circle (diameter: 11 mm) indicates the FOV of this system. (c) An image obtained by the combination of an sCMOS camera, Zyla 4.2, and the camera lens used in the AMATERAS1.0 system. The left panel shows the full FOV regulated by the image sensor, and the right two panels show the magnified views of the two local regions (*A*: center, *B*: corner).

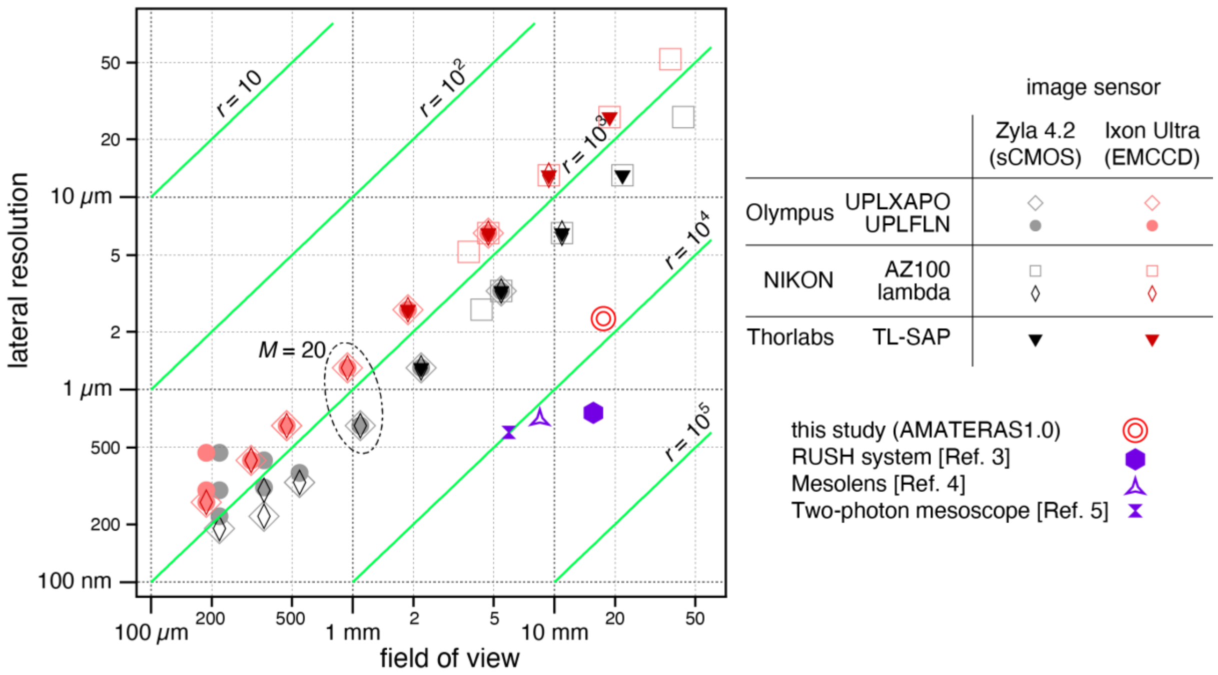


**Figure S3: Comparative plots of FOV and lateral resolution of imaging systems composed of conventional microscope objectives and image sensors.** Two series of Olympus, two series of NIKON, and a series of Thorlabs were selected as objective lenses, and Zyla 4.2 of Oxford Instruments were selected as image sensors. The contour lines of the trans-scalability index are shown in green color. The performance of this study (AMATERAS1.0) and three previously reported imaging systems (Refs. 3,4,5) are also plotted for the comparison of the lateral resolution corresponding to the FOV.

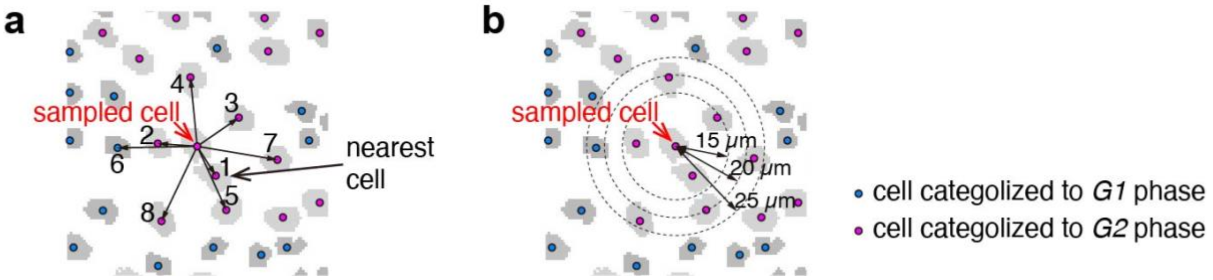


**Figure S4: Schematic illustration of the two analyses on similarity of neighboring cells.** (a-b) The rates of in-phase cells of neighboring cells as a function of (a) order of distance and (b) radius of neighboring area.

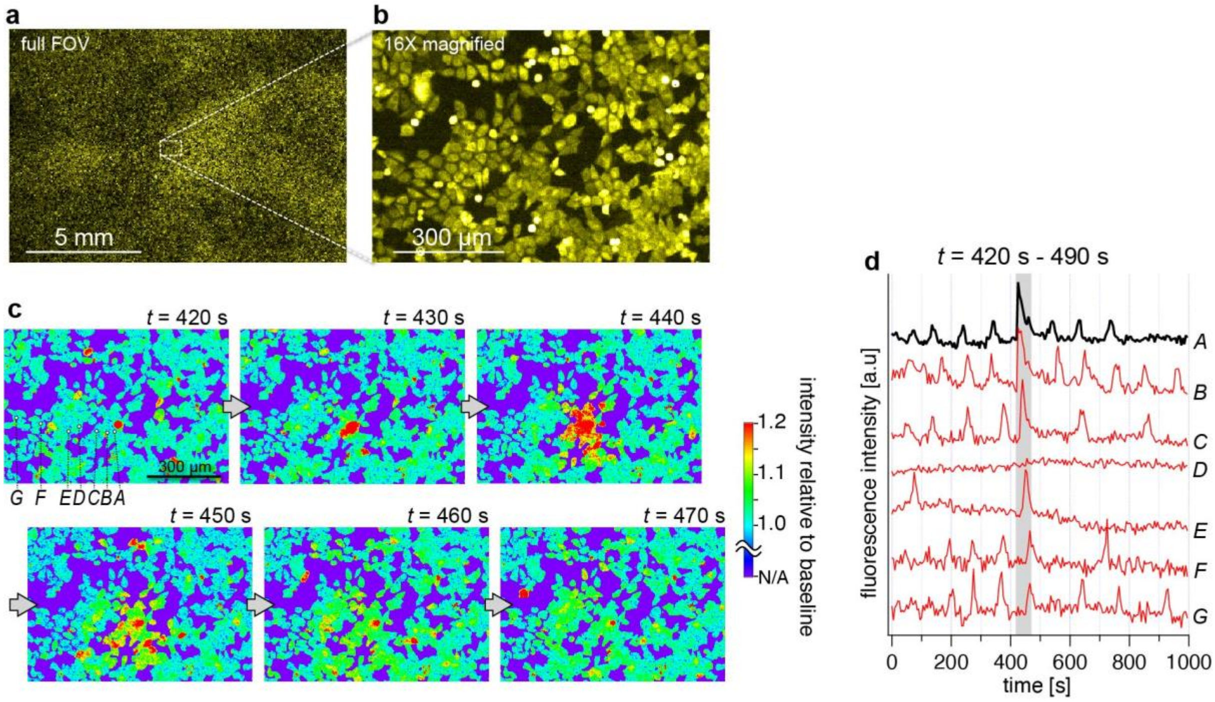


**Figure S5: Histamine-induced [$Ca^{2+}$] oscillation and spontaneous radial propagation observed in YC3.60-HeLa cells.** (a-b) Venus fluorescence images corresponding to (a) Full FOV image and (b) a 16X magnified view of the dashed rectangle indicated in (a). (c) Time course of the region indicated in (b) in 50 seconds of a particular time range ($t$ = 420 s - 490 s) where a radial propagation of [$Ca^{2+}$] wave was observed in this local region. The images are displayed with fluorescence intensity relative to baseline at each pixel. (d) Temporal profile of Venus fluorescence intensity of seven cells marked with *A*-*G* during the course of measurement (total time: 1000 s). The time range of the frames in (c) is indicated by the gray shade.

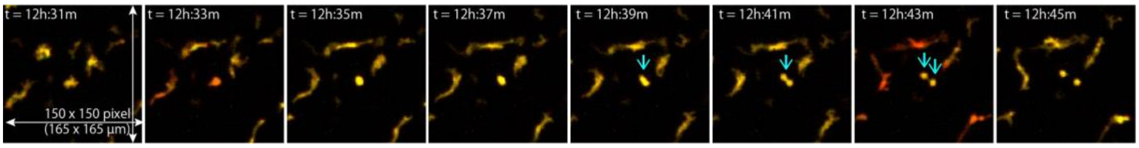


**Figure S6**: **A image sequence of a local region to show the dynamics of motion, morphology, and mitosis at the single-cell level**. Magnified views (165 μm × 165 μm) of a local region. A cell indicated by the downward arrow undergoes mitosis to be divided into two cells. It can also be seen that some other cells alter their morphology when moving around quickly. The imaging acquisition speed (30 s per frame) was high enough to track the single-cell movement along with changes in the morphology.
